# Use of a cytochrome P450 humanised mouse model to refine schistosomiasis drug discovery

**DOI:** 10.1101/2025.11.27.690949

**Authors:** Sarah D. Davey, Josephine E. Forde-Thomas, Benjamin J. Hulme, Kristin Lees, Alice H. Costain, Mary Evans, Gabriel Rinaldi, Laura Frame, Laste Stojanovski, Frederick R. C. Simeons, Amy Tavendale, A. Kenneth MacLeod, Remi Pichon, Yi-Hsuan Lee, Oktawia Polak, Iain W. Chalmers, Bismark Dankwa, Brenda Kisia Odhiambo, Victor Hugo Guimaraes, Matthew Hegarty, Martin T. Swain, Wayne Aubrey, Nicola Caldwell, Andrew S. MacDonald, Ian H. Gilbert, Beatriz Baragaña, Kevin D. Read, Karl F. Hoffmann

## Abstract

Control of schistosomiasis, a neglected tropical disease caused by infection with *Schistosoma spp*., remains reliant on a single chemotherapy, praziquantel (PZQ). This strategy presents a risk to global health should PZQ-resistant schistosomes establish in endemic areas and justifies the search for new drugs. However, species-specific metabolic differences between humans and preclinical models hinder the optimisation of next-generation anti-schistosomal therapeutics. Here, to bypass these species-specific limitations, we exploited a humanised mouse model, 8HUM, engineered to express the principal human Phase I cytochrome P450 enzymes (CYP1A1/2, CYP2C9, CYP2D6, CYP3A4/7) as well as the transcription factors constitutive androstane receptor (CAR) and pregnane X receptor (PXR) in place of 35 murine orthologues. We characterised *S. mansoni* development, immunopathology, hepatic transcriptomic responses, intestinal microbiome changes and PZQ metabolism as well as PZQ efficacy in 8HUM versus wild-type (WT) mice. 8HUM mice supported normal *S. mansoni* maturation, infection-associated microbiome dysbiosis, Th2-dominant immune responses and characteristic hepatic pathology. PZQ intrinsic clearance in 8HUM hepatic microsomes mirrored human levels and was >10-fold lower than that found for WT microsomes. Oral dosing revealed human-like PZQ exposures of (*R*)-PZQ and 4OH-PZQ in 8HUM mice at 25 mg/kg bodyweight and >90% reductions in worm burdens at 100 mg/kg bodyweight (equivalent to that seen in WT mice administered PZQ at 400 mg/kg bodyweight). Our results revealed that 8HUM mice recapitulate key features of murine schistosomiasis while exhibiting human-relevant drug metabolism. These findings establish 8HUM as a refined translational platform for anti-schistosomal drug development, improving predictive accuracy and accelerating therapeutic discovery.

**One Sentence Summary:** A cytochrome P450 humanised mouse model is used to study *Schistosoma mansoni* development, schistosomiasis, drug metabolism and drug efficacy.

## Introduction

Human schistosomiasis is a neglected tropical disease primarily caused by infection with three dioecious blood fluke species (*Schistosoma mansoni, Schistosoma haematobium* and *Schistosoma japonicum*). The disease mainly occurs in tropical and sub-tropical areas, kills thousands of individuals every year and contributes to an annual loss of up to 4.5 million disability adjusted life years (DALYs) in affected communities (*1*).

Over the last several decades, schistosomiasis has predominantly been controlled by praziquantel (PZQ), a drug that has recently been shown to activate both a transient receptor potential calcium channel in the parasite (SmTRPM_PZQ_) (*2, 3*) and a G-protein coupled receptor in the mammalian host (5HT_2B_) (*4*). While PZQ is relatively safe and inexpensive to administer, it does have documented limitations (*5*). One particular limitation includes significant first-pass metabolism in the liver leading to low systemic exposure in the bloodstream (*6*), which may be partially responsible for a lack of *in vivo* efficacy against juvenile (14-28 day old) parasites (*7*) and ultimately low cure rates in high-transmission areas (*8*). With preventative chemotherapy programmes distributing PZQ at an all-time high, there is also an increased fear that PZQ tolerant or resistant parasites could develop in endemic communities (*9, 10*). Collectively, these shortcomings support a rationale for identifying new anti-schistosomal compounds with mechanisms of action distinct from PZQ as a component of an integrated strategy to help advance the World Health Organisation goal of schistosomiasis elimination as a public health problem (*11, 12*).

A critical component of the schistosomiasis drug discovery pipeline is the use of preclinical animal models (e.g. mice) although, unavoidably, there are often major species differences in the metabolic routes of drug elimination. Metabolism of most approved small molecule therapeutics in humans is catalysed by the cytochrome P450 (CYP) superfamily (*13–16*). From the four CYP sub-families (CYP1A, CYP2C, CYP2D and CYP3A) involved in human xenobiotic metabolism, five produce enzymes responsible for greater than 90% of CYP-mediated drug metabolism (i.e. Phase I metabolism). In contrast, due to both environmental and evolutionary drivers, mice utilise a greater number of genes to catalyse similar Phase I processes (*17*). Therefore, there are inevitable differences in how rapidly these two species metabolise compounds/drugs. With mice being the dominant pre-clinical model for testing the *in vivo* efficacies of anti-schistosomal compounds, it is likely that mouse-specific metabolic processes have led to the attrition of many promising candidates.

To circumvent this challenge, an extensively humanised mouse line (8HUM) has recently been generated to reflect human Phase 1 drug metabolism pathways (*18*). This mouse line has had its 33 *cytochrome p450*s of the *cyp1a*, *cyp2c*, *cyp2d* and *cyp3a* gene subfamilies, together with the transcription factors *car (constitutive androstane receptor)* and *pxr (pregnane X receptor)* deleted and replaced with the major human Phase 1 drug metabolising *cytochrome P450* genes *CYP1A1, CYP1A2, CYP2C9, CYP2D6, CYP3A4* and *CYP3A7*, together with the transcription factors *CAR* and *PXR*. Recently, this model was successfully used to bypass mouse-specific metabolic limitations in efficacy models of *Mycobacterium tuberculosis*, *Leishmania donovani* and *Trypanosoma cruzi* infection (*19*), but has yet to be employed in the context of helminth infection.

By demonstrating in this study that 8HUM mice can fully support the sexual development of *S. mansoni*, are capable of mounting T helper cell type 2 (Th2) - dominant immunological and pathological responses to infection once egg laying occurs and can be used to quantify metabolism as well as drug-induced efficacy of PZQ, we contend that this model is well-positioned to drive a step-change advance in the schistosomiasis drug discovery pipeline. The use of this translational tool, to complement more traditional approaches in drug discovery, will circumvent mouse-specific metabolism disadvantages and allow optimisation of human pharmacokinetics (PK). This refinement has the potential to significantly reduce timelines and associated costs of progressing urgently-needed, new anti-schistosomal candidates.

## Results

### 8HUM mice support the development and sexual maturation of *Schistosoma mansoni*

An important pre-requisite in the use of 8HUM mice as a refined preclinical replacement to WT mice for early-stage schistosome drug discovery is the knowledge that these animals can support parasite development and mount Th2 dominant immunological and pathological responses to infection once egg laying commences. To assess these parameters, 80 cercariae were used to initiate percutaneous infection; at days 46-47 post-infection, parasitological and host immunopathological outputs as well as faecal microbiota compositions were compared between 8HUM and WT mouse strains (Fig. 1A). Infection did not negatively impact cumulative average weight gain when compared to uninfected controls; in fact, all mice (regardless of genotype or infection status) gained an average weight of ∼4-5 g (no significant differences between groups; *p* > 0.15) during the 47-days under study (Fig. 1B). While 8HUM mice harboured slightly fewer adult male and female schistosomes at the termination of the experiment (47 days post-infection), these worm burden differences were not significant (Fig. 1C, *p* = 0.72). To determine if mouse genotype affected total egg burdens or estimates of *in vivo* schistosome fecundity, eggs laid by adult females were isolated and counted from both liver and intestinal tissues of infected 8HUM and WT mice. Again, no significant differences were observed in the total amount of eggs per gram (EPG) of tissue (*p* = 0.96 for hepatic EPG; *p* = 0.18 for intestinal EPG) or when normalised by numbers of recovered females (*p* = 0.53 for hepatic comparison; *p* = 0.11 for intestinal comparisons; Fig. 1C). Furthermore, the sex-ratio normally seen in *S. mansoni* infected WT mice was also observed in 8HUM mice (8HUM mice, male: female ratio = 1.3; WT mice, male: female ratio = 1.6) (*20*).

**Fig. 1.**
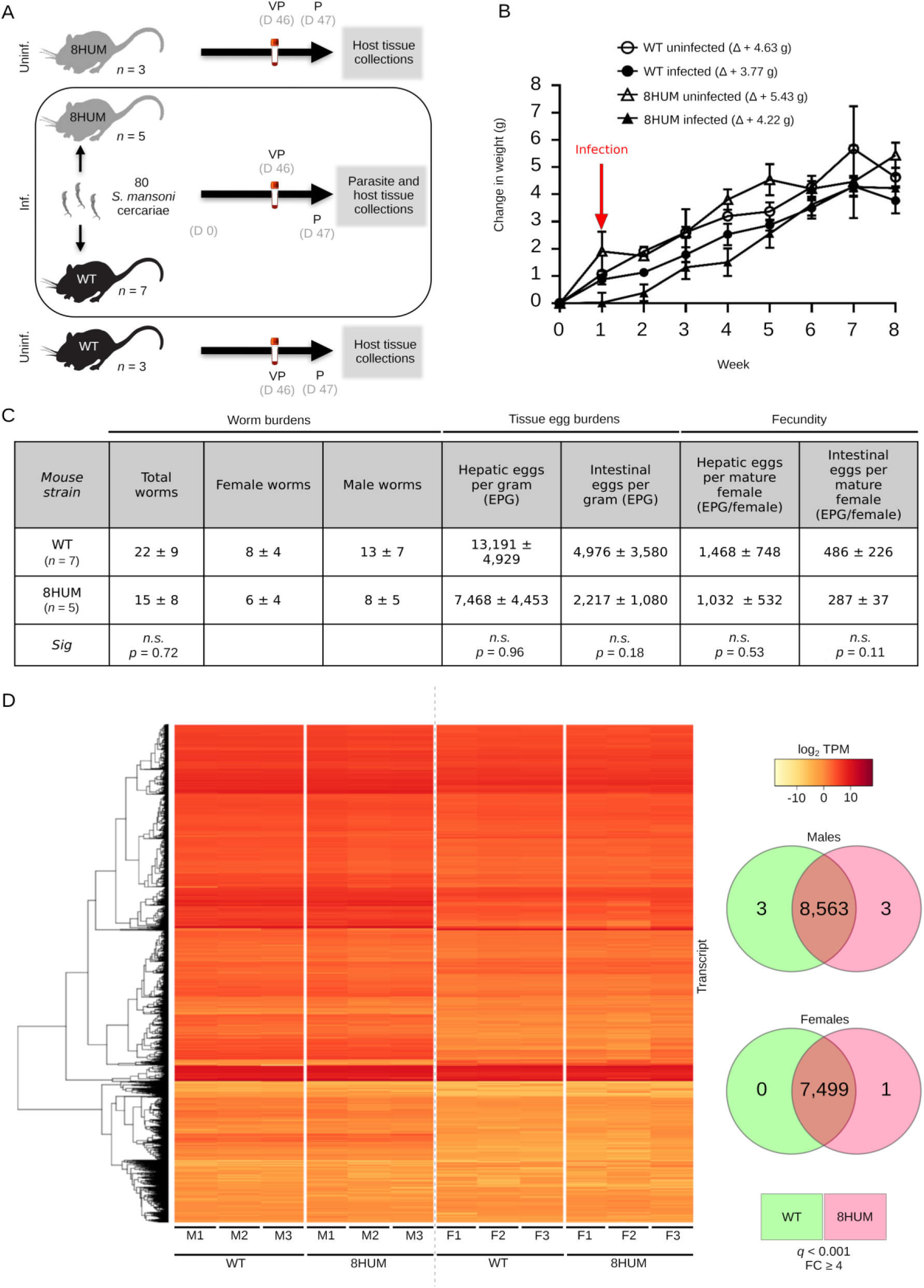
8HUM mice fully support the sexual development of *Schistosoma mansoni.* **(A)** Schematic overview of experimental process (VP: venous puncture, P: perfusion, Inf.: infected, Uninf: uninfected, D: day). **(B)** Average change in mouse weight from 0-8 weeks. Mice were infected with 80 cercariae and weight change was recorded weekly, starting one-week prior to infection (week 0). The change in weight was expressed as the mean weight± SEM at each time point subtracted from the initial weight at week 0. **(C)** Summary of worm counts, tissue egg burdens and fecundity following infection with 80 cercariae. **(D)** Transcript expression (log_2_ transcripts per million, TPM) in male and female worms perfused from WT or SHUM mice 47 days post-infection. VENN diagrams to the right of the heatmap represent ditterential expression at q < 0.001 and FC ≥ 4, with shared (non-DE) genes in the overlapping sections (8,563 genes in males and 7,499 genes in females) and significantly up-regulated genes for each strain in non-overlapping sections.

Previous studies have shown that liver microsomes obtained from 8HUM mice metabolise drugs quite differently than liver microsomes derived from WT mice (*19*). Therefore, differential liver metabolism in infected 8HUM animals could lead to the production of metabolites that impact the development of adult worms residing in the hepatic portal system. To indirectly assess this hypothesis, we analysed the transcriptomes of recovered worms by bulk RNA-Seq (Fig. 1D). Here, adult male and female transcriptomes were compared from schistosomes obtained from both 8HUM (*n* = 3) and WT (*n* = 3) mice using stringent *q* value thresholds (*q* < 0.001) for identifying differentially expressed transcripts (FC ≥ 4 or log_2_FC ≥ 2) (Fig. 1D, Data file S1). From 9,890 predicted *S. mansoni* protein coding genes (*smps*) present in the *S. mansoni* genome assembly (v.10) (Fig. 1D, heatmap), a total of 8,563 *smps* were expressed (i.e. transcripts with any detectable expression > 0) in males regardless of the mouse strain in which parasites developed. Only six *smps* were differentially expressed (three up-regulated in males derived from 8HUM mice and three up-regulated in males derived from WT mice; Fig. 1D, Venn diagram and Data file S2). Indeed, male transcriptomes were remarkably similar (Pearson correlations presented in Fig. S1). This trend was mirrored in female schistosomes where 7,499 *smps* were expressed (i.e. transcripts with any detectable expression > 0) and only one *smp* was differentially expressed in females derived from 8HUM mice (Data file S2 and Fig. S1, Venn diagram). Even when *q* value thresholds were relaxed (*q* < 0.05), only one additional *smp* was identified as being differentially expressed in male transcriptomes (*smp_093430.2;* highlighted row, Data file S2); no further differentially expressed *smps* were identified in female transcriptomes (Data file S2). These parasitological and transcriptomic data indicate that extensive humanisation of the cytochrome P450 pathway did not significantly impact *S. mansoni* development.

### Immunological responses in 8HUM mice are comparable to WT animals

Having established that 8HUM mice are susceptible hosts for *S. mansoni*, we next assessed whether they immunologically responded to infection like WT mice (Fig. 2). Serum, obtained from blood samples at day 46 post-infection, demonstrated anti-soluble worm antigen preparation (SWAP) IgG (total), IgG_2b_ and IgG_1_ responses that were slightly higher (not significant; *p* > 0.90 for all isotypes) in 8HUM mice when compared to WT animals (Fig. 2A). Anti-soluble egg antigen (SEA) IgG (total) and IgG_2b_ responses were also slightly higher in infected 8HUM mice compared to infected WT animals (again not significant; *p* = 0.87 and *p* = 0.71 respectively). However, infected 8HUM mice mounted an approximate two-fold increase in anti-IgG_1_ SEA that was statistically different than that measured in infected WT animals (*p* = 0.01).

**Fig. 2.**
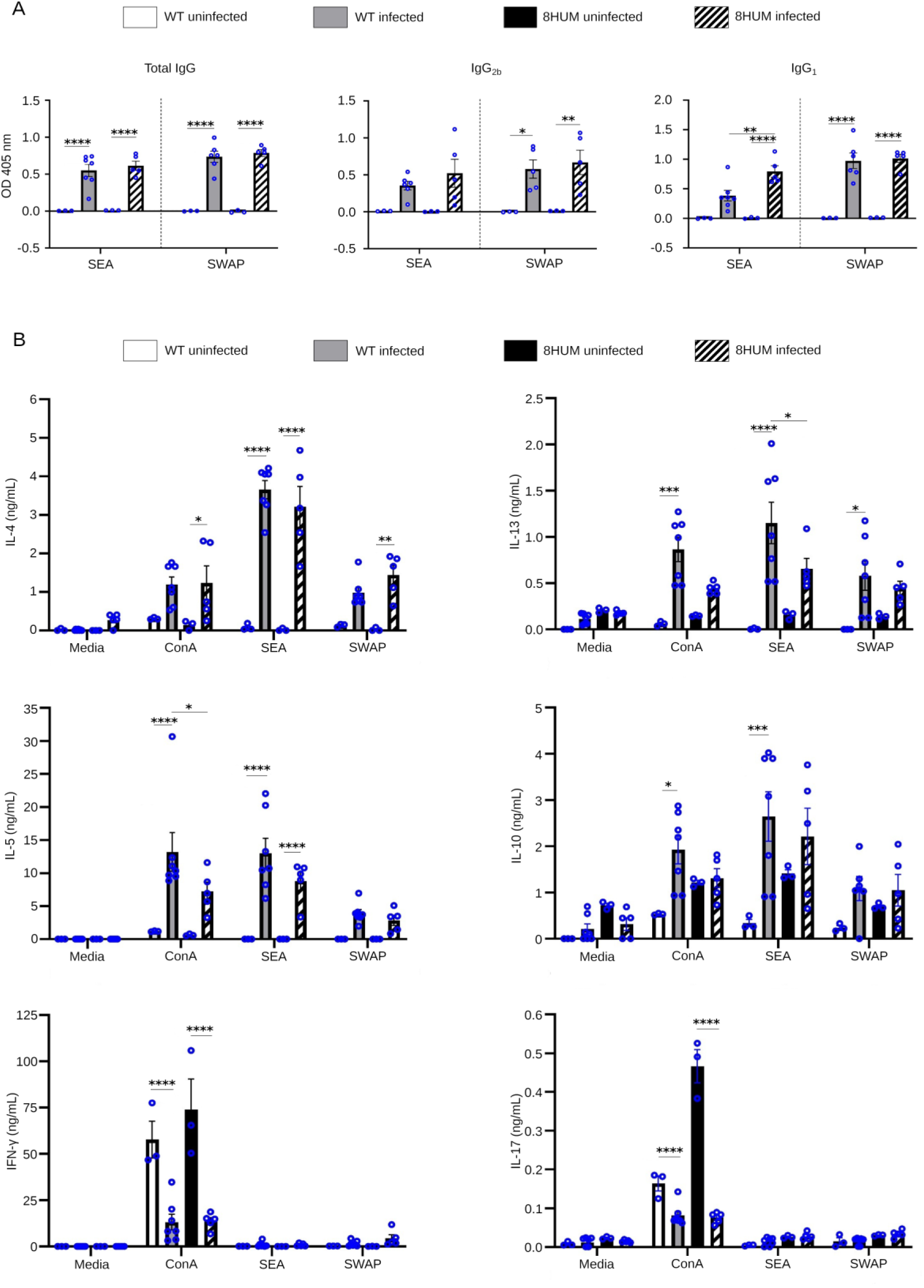
Immunological responses in SHUM mice are comparable to WT mice. **(A)** Antigen specific lgG antibody responses (Day 46) from mouse sera and (B) antigen specific/ConA stimulated cytokine responses of splenocyte cultures (Day 47) were assessed by ELISA. For both, data is expressed as mean ± SEM. Statistically significant differences were calculated using a two-way ANOVA with Tukey’s Multiple Comparison Test. Differences within mouse species and between the two infected strains are shown with an asterisk where • *p* < 0.05, •• *p* < 0.01, ••• *p* < 0.001, and •••• *p* < 0.0001 *(n* = 3 for *WT* uninfected and BHUM uninfected, *n* = 7 for *WT* infected and *n* = 5 for BHUM intected). Samples have been background corrected.

To compare anti-schistosomal cellular responses to infection, splenocytes derived from 8HUM and WT spleens at day 47 post-infection were re-stimulated with SWAP and SEA (Fig. 2B). Th1 (IFN-gamma), Th2 (IL-4, IL-5, IL-13), anti-inflammatory (IL-10) and Th17 (IL-17) cytokines were measured in the culture supernatants at 72 h post-stimulation. SWAP and SEA-specific splenic production of Th2 cytokines (IL-4, IL-5 and IL-13) and anti-inflammatory IL-10 were all elevated upon schistosome infection in both 8HUM and WT mice. In contrast, very little IL-17 or IFN-gamma was produced in response to antigen (or Concanavalin A; ConA) stimulation in infected 8HUM or WT splenocyte cultures despite both cytokines being detectable in uninfected splenocyte supernatants stimulated by ConA. SEA induced IL-13 (as well as ConA stimulated IL-5) was the only antigen-specific cytokine that was statistically reduced in 8HUM splenocyte cultures when compared to WT samples (*p* = 0.02). Taking both antigen-specific antibody and splenic cytokine responses together, these data suggest that, although there were some differences in baseline levels, 8HUM mice immunologically-responded to *S. mansoni* infection similarly to WT mice.

### Infection-associated faecal microbiota dysbiosis occurs in 8HUM mice

*S. mansoni* infection in WT mice affects host microbiota dysbiosis (i.e. reduced alpha diversity) in faecal matter collected from intestines (*21, 22*). To understand if this infection-associated process develops similarly in 8HUM mice, 16S rRNA sequencing of microbial DNA isolated from intestinal contents was compared between 8HUM and WT mouse groups (uninfected and 47 days post-infection; Fig. S2). As previously reported (*21, 22*), *S. mansoni* infection of WT mice led to a decrease in alpha diversity of faecal microbiota communities (Fig. S2A). A similar trend was also observed in 8HUM faecal samples, indicating that infection of both mouse strains was linked to a reduction in bacterial species richness in intestinal contents. However, differences in species composition (i.e. beta diversity) were predominantly driven by mouse genotype, regardless of infection status (Fig. S2B). Collectively, these observations are entirely consistent with previous reports studying schistosome-associated effects on microbiota in different mouse strains (*21, 22*) and suggest that extensive humanisation of the cytochrome P450 pathway is not a confounder.

### Infection-induced hepatic transcriptomes are minimally affected by cytochrome P450 humanisation

*S. mansoni* infection in WT mice leads to transcriptional changes in hepatic genes responsible for inflammation, extracellular matrix remodelling and wound healing (*23, 24*). To quantify how these pathology-associated features develop in 8HUM mice, bulk RNA-Seq of liver tissues was compared between WT and 8HUM animals (uninfected and 47 days post-infection groups; Fig. 3). As the 8HUM mice should be deficient in 33 transcripts derived from four *cyp450* gene subfamilies (*cyp1a*, *cyp2c*, *cyp2d and cyp3a*) as well as the transcription factors *car* and *pxr* (*18*), we first examined liver transcriptomes from highly correlated replicates (*n* = 3 for uninfected samples, *n* = 4 for infected samples; Fig. S3) for evidence of these products’ expression (Fig. 3A, Data file S3). Unsurprisingly, and regardless of infection status, 8HUM transcriptomes had negligible expression (average TPM ≤ 2) for most mouse *cyp450* subfamilies (including two *cyp3a41* paralogs, *cyp3a41a* and *cyp3a41b*). Five genes (*car*, *cyp1a2*, *cyp2c70*, *cyp3a57* and *pxr*) were still detectable (maximum TPM > 10) in 8HUM hepatic transcriptomes, however, all cases were explained by residual mappings to incomplete transcripts (e.g. retention of non-functional exons 6-9 in *cyp2c70*) or UTR sequence (Fig. S4). In WT mice, schistosome infection led to a reduction in most members of these cytochrome *p450* gene subfamilies and transcription factors; *cyp2c70* was the single exception and was significantly induced upon infection (average TPM counts = 488.24 in uninfected animals; average TPM counts = 3,759.23 in infected animals, *q* < 0.001, log_2_FC ≤ 0.530).

**Fig. 3.**
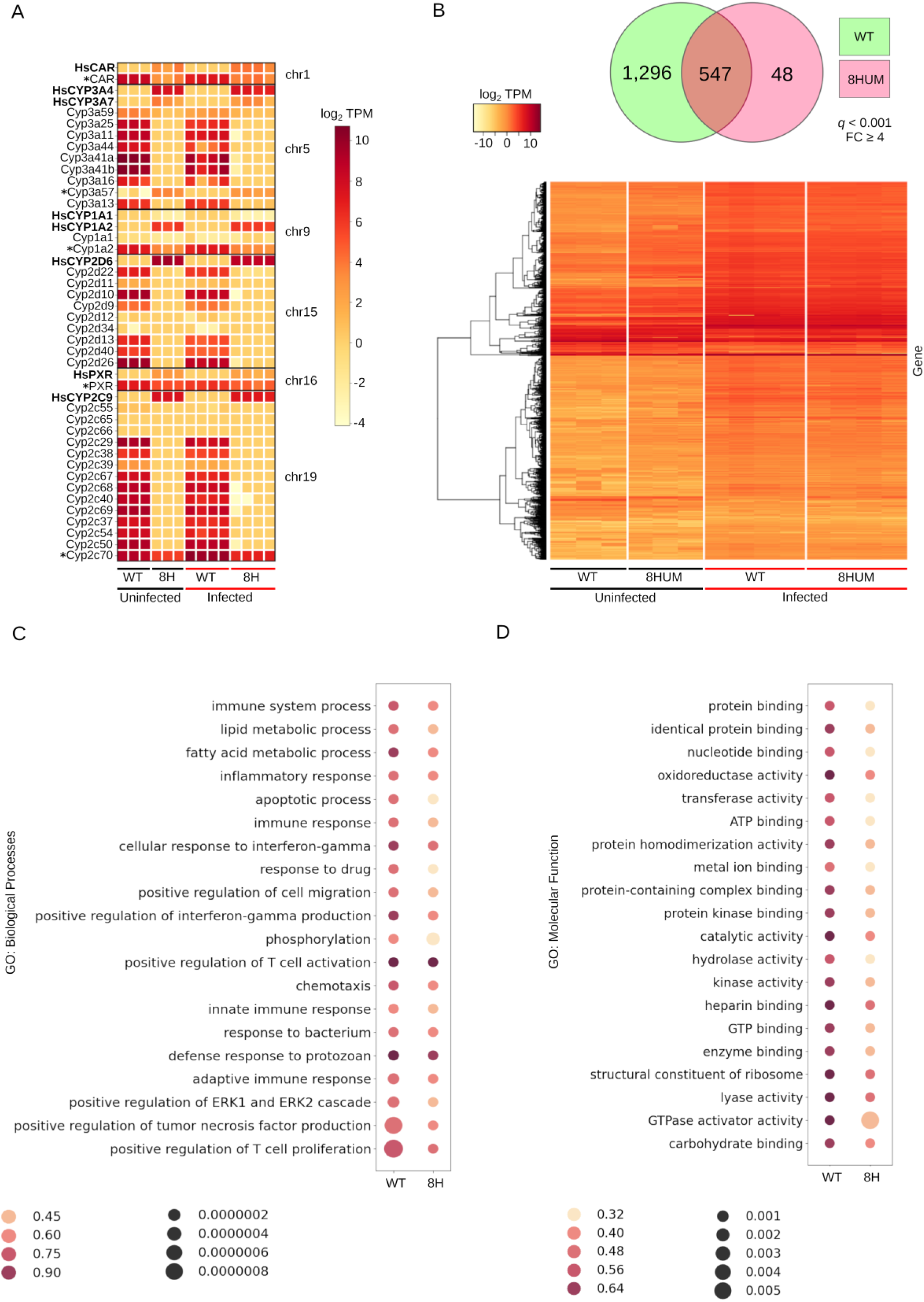
Hepatic transcriptomes in SHUM mice respond to S. *mansoni* infection similarly to *wr* mice. **(A)** Expression heatmaps (log_2_ transcripts per million, TPM) of the 35 *cytochrome p450* genes and transcription factors *car* and *pxr* substituted for human orthologues in SHUM mice *(cyp3a41* has two paralogs: *cyp3a41a* and *cyp3a4lb*) in addition to their 8 humanised counterparts. * indicate mouse genes with residual transcription in SHUM (max TPM >10). **(B)** Expression (log2 TPM) of differentially expressed (DE) genes (q < 0.001 and FC ≥ 4) between uninfected and infected (at D 47 post-infection) *wr* and SHUM hepatic transcriptomes. The VENN diagram above the heatmap indicates distinct and shared genes which are DE in response to infection. *wr* and SHUM mice differentially expressed 1.296 and 48 unique genes, respectively. A total of 547 DE genes in response to infection are found in common. **(C)** Top twenty enriched GO terms for gene sets within Biological Processes and **(D)** Molecular Function in response to infection. In both bubbleplots **(C** and **D),** enrichment ratio (proportion of genes annotated by a term that are significantly enriched where 1 = all oenes enriched) is shown bv dot colour and sionificance is shown bv dot size (smaller dots = more sionificant, smaller a value).

Expression of knocked-in human genes was also detected in all 8HUM samples, with *CYP2C9* most highly expressed (average TPM 695.46 for uninfected mice and 398.68 for infected mice, Data file S3). Most 8HUM humanised cytochromes decreased in expression in response to infection, although this decrease was only significant (*q* < 0.05) in two instances: *CYP2C9* (*q* < 0.01, log_2_FC = 0.451) and *CYP3A4* (*q* < 0.05, log_2_FC = 0.664). Transcription factor *CAR* was the exception, marginally increasing in response to infection (from an average TPM of 8.899 to 14.232); however, this change was not significant (*q* = 0.436, log_2_FC = 0.370). Next, we examined how schistosome infection/egg embolisation globally affected transcription in the liver. From a total of 17,572 *M. musculus* protein coding genes (genome assembly GRCm39) with detectable expression > 0 (Data file S4), we quantified hepatic transcripts that were differentially expressed (up- or down-regulated) upon infection in each mouse strain using a stringent *q* value cutoff of 0.001 and a FC ≥ 4 (Fig. 3B, Data file S5).

Using these criteria, we found that infection led to the differential expression of 1,891 genes (heat map, Fig. 3B). When deconvoluting these results according to mouse strain, infection in WT mice led to the differential expression of 1,296 unique liver transcripts whereas infection in 8HUM animals led to the differential expression of 48 unique liver transcripts; an overlapping set of 547 transcripts were differentially expressed upon infection in both mouse groups (Venn diagram, Fig. 3B). Gene set enrichment analyses (GSEA) revealed a top twenty list of shared Gene Ontology (GO) Biological Processes (BP) and Molecular Function (MF) terms that were significantly over-represented upon infection (*q* < 0.05) in both 8HUM and WT mice (Figs. 3C and 3D, respectively). As illustrated in the bubbleplots, the numbers of enriched genes within the top twenty term sets were higher in response to infection in all BP (Data file S6) and MF (Data file S7) terms in WT mice compared to 8HUM mice. This GSEA demonstrated that both mouse strains mounted a qualitatively similar transcriptional response to schistosome infection/egg embolisation in the liver, though, quantitatively the proportion of differentially expressed genes within each gene set varied between strains.

### Characteristic hepatic pathology develops in infected 8HUM mice

*S. mansoni* infection in WT mice leads to the formation of collagen-enriched granulomas around eggs trapped in an inflamed liver (as reviewed in (*25*)). Upon gross pathological examination of murine pathology at necropsy, schistosome infection clearly led to similar, egg-induced macroscopic lesions in enlarged 8HUM livers (Fig. 4A). Histological examination of these lesions clearly revealed defined (haematoxylin and eosin; H&E) and collagen-rich (picrosirius red; PSR) granulomas (Fig. 4B). However, these hallmarks of murine schistosomiasis were slightly larger (*p* < 0.01) and more collagen-enriched (*p* = 0.09) in WT mice (Fig. 4C). The modest increase in granulomatous pathology observed in WT mice correlated with similar increases in liver *il-13* (*q* < 0.001, log_2_FC = 1.39) and *il-4* (*q* = 0.15, log_2_FC = 0.81) (Fig. 4C), two cytokine gene products responsible for positively regulating granuloma formation and hepatic fibrosis during murine schistosomiasis (*26*). Further GO analyses of BP terms involved in fibrosis and granuloma formation (collagen fibril organisation (GO:0030199), Data file S8; and extracellular matrix organisation (GO:0030198), Data file S9) provided additional support for the hepatic pathology differences observed between infected WT and 8HUM mice (Fig. 4D). Here, a subset of genes in each BP term was significantly up-regulated in response to infection in both WT and 8HUM animals (Fig. 4D, Both); however, a larger subset was significantly increased upon infection in WT mice only (Fig. 4D, WT only). Many of these genes have previously been positively linked to granuloma formation and hepatic fibrosis during murine schistosomiasis (*23, 24, 27–29*); their greater induction in WT mice upon infection provides an explanation for the modest increases in granuloma volume and intra-granuloma fibrosis observed in our study. However, despite differences in magnitude, these histopathological and transcriptional findings collectively indicated that 8HUM mice develop the characteristic Th2-driven, pathological features of experimental schistosomiasis seen in WT animals.

**Fig. 4.**
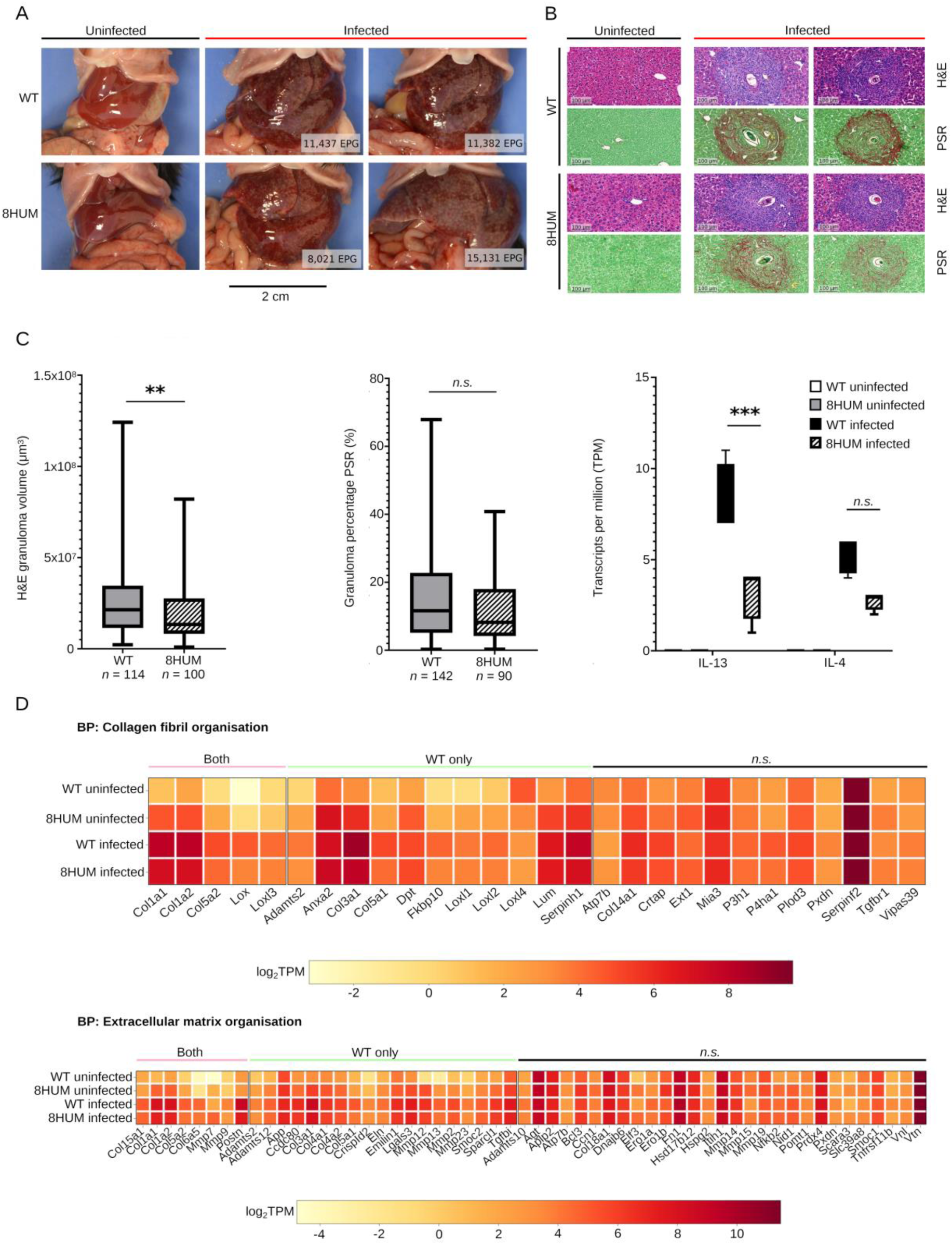
SHUM mice develop characteristic granulomatous hepatic pathology upon S. *mansoni* infection. **(A)** Representative necropsy images from livers of uninfected *(n* = 1) and infected (n = 2) WT and SHUM mice. Average EPGs of lrver tissue for representatrve infected mdiv1duals are provided in white boxes. **(B)** Uninfected *(n* = 1) and infected *(n* = 2) mouse livers stained with haematoxylin and eosin (H&E) and picrosirius red (PSR). **(C)** Quantification of liver pathology-associated parameters include: granuloma volumes calculated from H&E areas assuming perfect sphericity *(p* < 0.01 according to Mann-Whitney u test); percentage collagen content per granuloma as measured by quantification of PSR+tissue *(p* = 0.09 according to Manr1-Whitney U test); infection-induced increases in expression of cytokines *il-13* (q < 0.001, log,FC = 1.39) and *il-4* (q = 0.15, log,FC = 0.81). **(D)** Expression heatmaps (log, TPM) of genes annotated by GO terms collagen fibril organisation (GO:0030199) and extracellular matrix organisation (GO:0030198) with > 10 TPM across all replicates. Genes have been categorised into either ’Both’, ’*WT* only’ or ’n.s.’ to indicate significant differentialexpres.sion in both *WT* and SHUM, significant differential expressionin WT only or no significant differential expression in response to inl’ection, respectively.

### Translational relevance of PZQ metabolism is improved in the 8HUM model

To investigate the suitability of the 8HUM model for assessing drug-induced efficacy in schistosome-infected animals, we first explored the *in vitro* clearance rate (intrinsic clearance; CL_int_) of PZQ by 8HUM hepatic microsomes and *in vivo* pharmacokinetics (PK) of PZQ in uninfected 8HUM mice (Fig. 5). *In vitro*, 8HUM and human hepatic microsomes metabolised racemic PZQ (rac-PZQ) at a similar rate (8HUM CL_int_ = 2.7 mL/min/g, human CL_int_ = 3.2 mL/min/g); noticeably, these CL_int_ values were > 10-fold lower than that measured for WT mouse hepatic microsomes (CL_int_ = 37.0 mL/min/g) (Fig. 5A). *In vivo*, as measures of exposure (AUC and C_max_) in 8HUM mice can be far higher than in WT mice administered the same dose (*19*), we conservatively orally-dosed 8HUM animals with 25 mg/kg rac-PZQ compared to 400 mg/kg rac-PZQ in WT mice for an initial range-finding PK study (Fig. 5B). Both groups of treated animals displayed similar PK profiles for the active enantiomer of PZQ (i.e. *R*-PZQ) and the major metabolite of PZQ (4OH-PZQ) throughout the study (Fig. 5C). Despite a 16-fold difference in administered dose, AUC_last_ and C_max_ values of *R*-PZQ in 8HUM animals were only ∼4-fold lower when compared to WT animals; AUC_last_ and C_max_ values for 4OH-PZQ in 8HUM animals were ∼5-fold lower (Fig. 5D). Based on these exposure differences, and assuming a linear relationship between dose and exposure, we predicted that oral administration of a single dose of ∼100-200 mg/kg rac-PZQ to *S. mansoni*-infected 8HUM animals would lead to similar levels of parent drug and metabolite exposure and, thereby, drug-induced efficacies (∼94%) comparable to that typically observed in WT animals receiving a single oral dose of 400 mg/kg rac-PZQ (*30*).

**Fig. 5.**
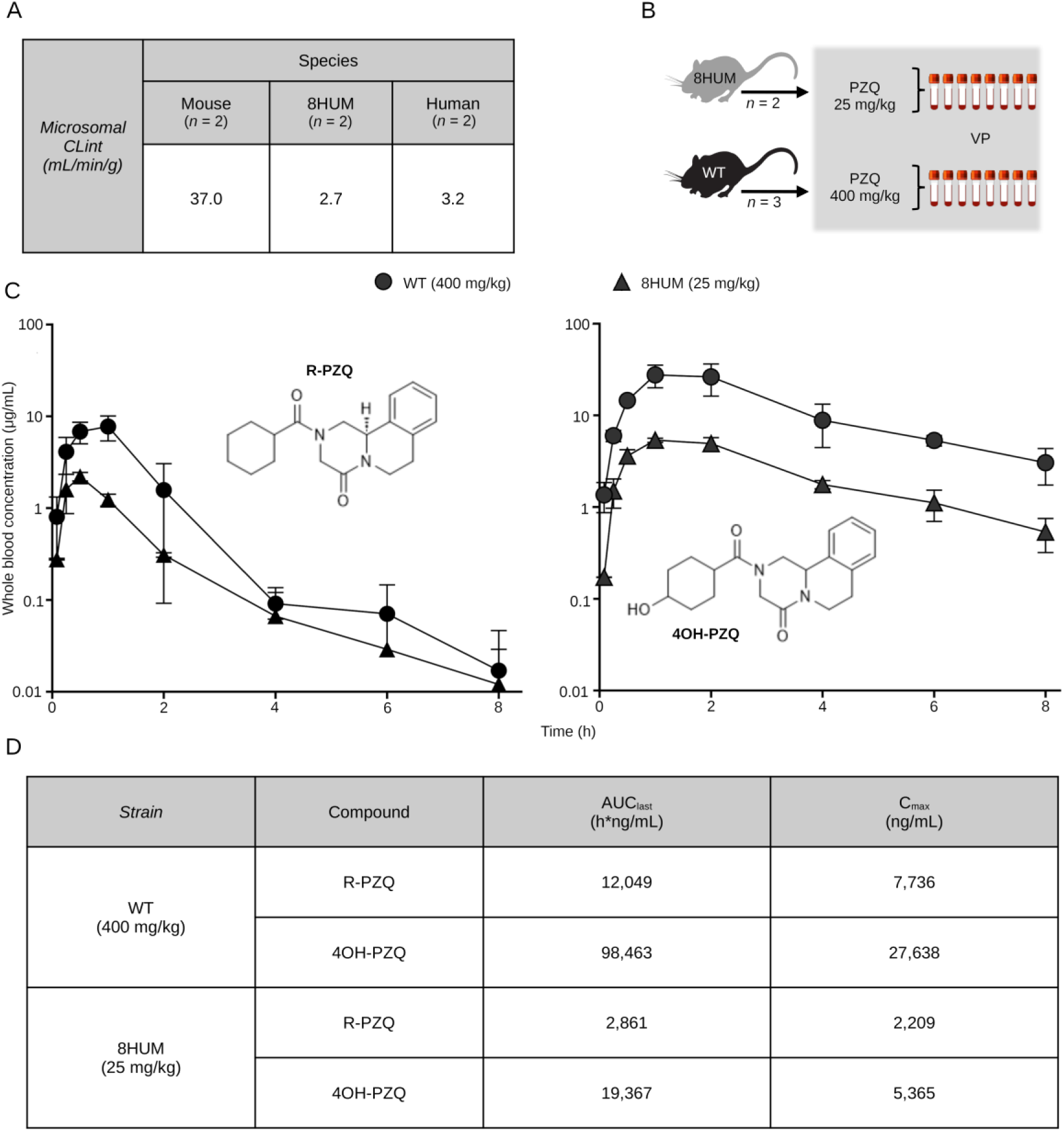
8HUM mice exhibit human-equivalent PZQ metabolism. **(A)** In vitro clearance rate of PZQ by liver microsomes (n = 2 for each species; intrinsic clearance: CLint). **(B)** Schematic overview of range-finding pharmacokinetic study. Blood samples were obtained by venepuncture (VP) at 5,15, 30, 60, 120, 240, 360, and 480 min after rac-PZQ administration. **(C)**. Concentration versus time plots of total blood levels are shown for R-PZQ and R-Trans-4OH-PZQ for both WT (n = 3; 400 mg/kg) and 8HUM (n = 2; 25 mg/kg) animals following oral administration of rac-PZQ (mean concentrations ± SD shown). **(D)** Summary PK exposure levels for R-PZQ and R-Trans-4OH-PZQ in rac-PZQ dosed WT and 8HUM mice.

### Validation of the 8HUM model for quantifying PZQ-induced efficacy against *S. mansoni*

To test this prediction, groups of WT and 8HUM mice were infected with 120 *S. mansoni* cercariae and the efficacy of rac-PZQ treatment in differentially dosed groups was compared (Fig. 6). Specifically, at 41 days post-infection, 8HUM animals were orally dosed with excipient only (0 mg/kg; *n* = 10), 100 mg/kg (*n* = 10) or 200 mg/kg (*n* = 10) rac-PZQ. In comparison, WT animals were orally dosed with excipient only (0 mg/kg; *n* = 9) or with 400 mg/kg (*n* = 9) rac-PZQ (Fig. 6A). At day 47 post-infection, worm and egg burdens were compared between all groups (Fig. 6B). As previously observed (*30*), a single 400 mg/kg dose of rac-PZQ in WT mice led to a significant reduction (93%) in adult worm burdens (females (95%) > males (90%)) when compared to excipient controls. While a similar worm burden reduction (92%) was observed in 8HUM mice receiving a single 100 mg/kg dose of rac-PZQ, a greater worm burden reduction (98%) was found in the 8HUM group receiving a single 200 mg/kg dose of rac-PZQ (*p* > 0.99). Comparable to treated WT animals, rac-PZQ had a larger effect on female (vs male) worm burdens in both 8HUM treated groups. The effect on liver and intestinal egg burdens broadly mirrored worm burden differences found amongst all rac-PZQ treated groups.

**Fig. 6.**
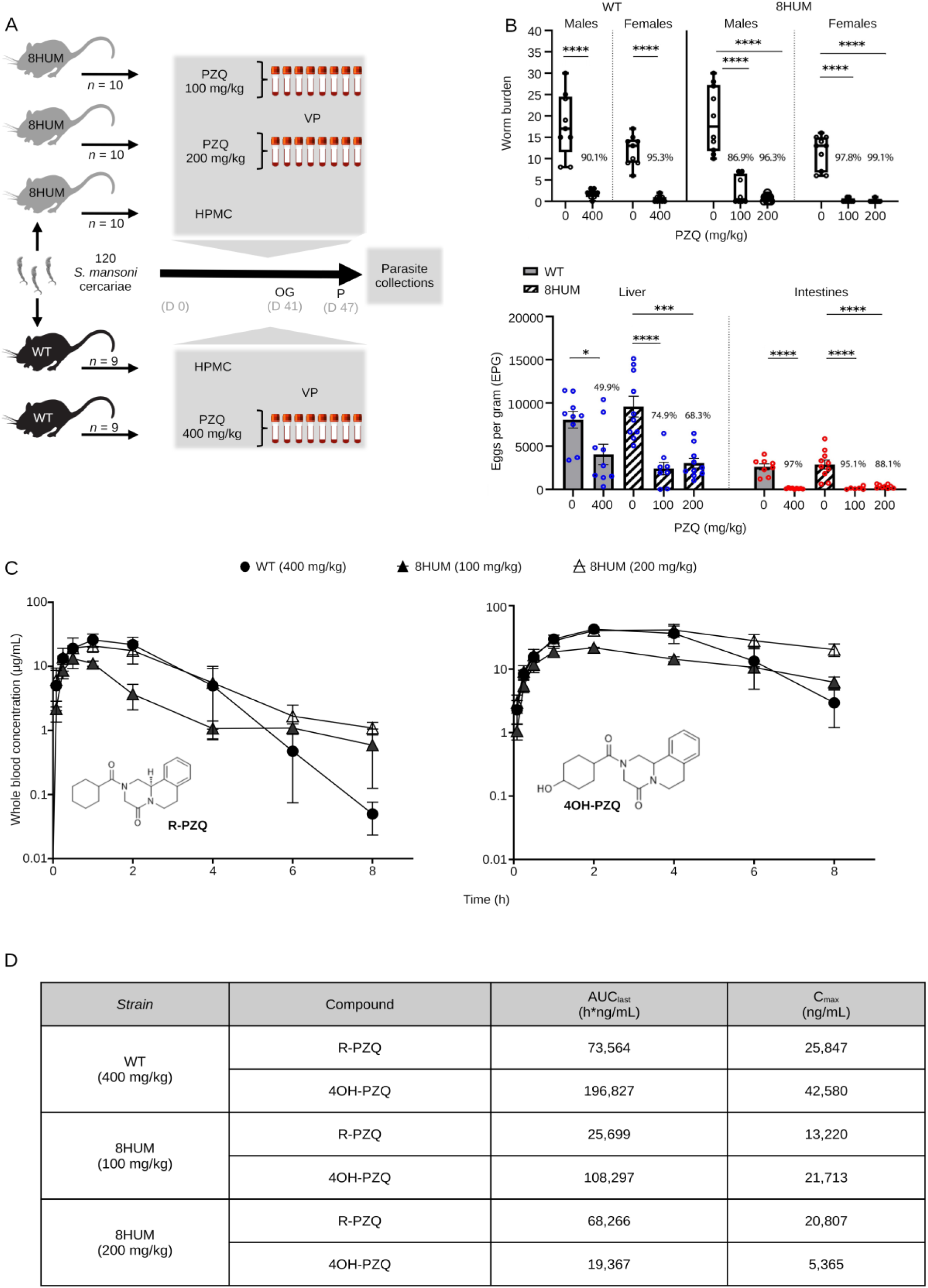
SHUM mice are an improvement over WT animals in pre-clinical efficacy studies. **(A)** Schematic overview for quantifying PZQ-induced efficacy in S. *mansoni* infected mice (VP: venous puncture, P: perfusion, OG: oral gavage, HPMC: hydroxypropyl methylcellulose (vehicle control), D: day). **(B)** The percent worm and egg reductions following exposure to PZQ is indicated above each of the groups and was calculated according to the formula: (untreated mean - PZQ treated mean/untreated mean x 100). Statistically significant differences in worm and egg burdens between untreated and PZQ-treated groups were determined by a one-way ANOVA with Tukey’s Multiple Comparison Test where* *p <* 0.05, *** *p* < 0.001 and ott *p* < 0.0001. **(C)** Concentration versus time plots of total blood levels are shown for R·PZQ and R-Trans-4OH-PZQ for both WT *(n =* 3, 400 mg/kg) and BHUM *(n =* 3, 100 mg/kg; *n =* 3, 200 mg/kg) animals following oral administration of rac-PZQ (mean concentrations ± SD shown). **(D)** Summary PK exposure levels for R-PZQ and R-Trans-4OH-PZQ in rac-PZQ dosed WT and BHUM mice.

PK analyses of blood samples derived from rac-PZQ-treated, infected animals demonstrated a ‘flattening out’ effect for both *R*-PZQ and 4OH-PZQ curves in the 8HUM groups compared to the WT group (Fig. 6C). This phenotype has been observed before in 8HUM animals dosed with a variety of structurally diverse, anti-infective medicines and may be driven by slower drug/compound elimination (*19*). Exposure (i.e. AUC_last_ and C_max_ values) of *R*-PZQ and 4OH-PZQ in infected 8HUM animals tracked alongside dosage administered and, for the 200 mg/kg treatment group, approached or exceeded the levels observed in infected WT mice dosed with 400 mg/kg (Fig. 6D). These results support the accuracy of our estimated rac-PZQ dosing range (100–200 mg/kg) required to elicit worm and egg burden reductions in infected 8HUM mice comparable to those observed in infected WT mice administered 400 mg/kg rac-PZQ.

## Discussion

Schistosomiasis is a neglected tropical disease predominantly managed by monotherapy with praziquantel (PZQ). Considering that there are no vaccines near clinical use (*31*), there are potential PZQ-resistant schistosome populations already established in high transmission areas of East Africa (*10*) and there is only one advanced PZQ replacement candidate under investigation as a late lead (*32*), the development of new treatments for schistosomiasis remains a public-health priority. To accelerate the progression of actionable new anti-schistosomal therapeutics, we comprehensively detail how the most advanced genetically-humanised mouse model for studying Phase I metabolism (8HUM (*18*)) can be successfully used as an integral component of current schistosome drug discovery pipelines.

As 8HUM mice have not previously been tested for their susceptibility to *S. mansoni* infection, nor have they been formally assessed for PZQ exposure during PK investigations, we sequentially conducted our study in three distinct phases to mitigate any adverse events. The first phase involved a robust characterisation of the salient features of schistosome infection in this mouse model and quantitatively assessed parasite development, host responses to infection and intestinal microbiota dysbiosis. When contrasted to infection in WT mice, most outputs measured in infected 8HUM mice were comparable. Nevertheless, slight differences in hepatic pathology (granuloma volume, intra-granuloma fibrosis and genes associated with both collagen fibril- and extracellular matrix-organisation) were observed. While not always significant, these differences were correlated with IL-13 levels (splenic-derived protein: Fig. 2, liver-derived transcript: Fig. 4). As parasite-induced IL-13 drives hepatic pathology during murine schistosomiasis (*26*), the slight decrease in parasitological load found in infected 8HUM mice could, in part, explain these liver alterations. However, metabolites differentially created in response to the cytochrome P450 repertoire of infected 8HUM animals, could also contribute to these observations. In support of this contention, a recent study concluded that 8HUM hepatic microsomes incubated with 14 approved human medicines created products that are more comparable to human than WT mouse metabolites (*19*). Therefore, global metabolomic profiling, in response to *S. mansoni* infection, is currently being performed in 8HUM mice. These metabolites might also contribute to the slight, but significant, differences observed in adult schistosome transcriptomes derived from 8HUM mice (6 differentially expressed transcripts in adult males and 1 in adult females). Nevertheless, our comprehensive characterisation of *S. mansoni* development and schistosomiasis-related changes in intestinal microbiota, immunology and hepatic immuno-pathology clearly indicates that 8HUM mice are suitable replacements for WT mice in the schistosome drug discovery pipeline.

The second phase of our studies involved the *in vivo* (PK) and *in vitro* (CL_int_) characterisation of rac-PZQ metabolism in uninfected 8HUM mice. The finding that 25 mg/kg bodyweight rac-PZQ administered to 8HUM mice led to systemic exposures of *R*-PZQ (AUC_last_ = 2,861 h*ng/mL) similar to those found in humans given 20-40 mg/kg bodyweight rac-PZQ (AUC_last_ = 1,303 – 4,830 h*ng/mL; (*33, 34*)) clearly indicates comparable rates of drug metabolism. 4OH-PZQ (both *R* and *S* enantiomers) systemic exposures in 8HUM mice (AUC_last_ = 19,367 h*ng/mL) also was within the range found for (*R*)-4OH-PZQ in human (children) populations given 20-40 mg/kg bodyweight rac-PZQ (AUC_last_ = 16,840 - 56,590 h*ng/mL) (summarised in (*35*)). Similarly, both 8HUM- and human-hepatic microsomes cleared rac-PZQ at near identical rates (CL_int_ values 2.7 mL/min/g vs 3.2 mL/min/g, respectively), which was substantially lower than that measured for WT-hepatic microsomes (CL_int_ = 37 mL/min/g). Thus, the removal of high activity mouse enzymes that limit exposure to PZQ highlights a key advantage of using 8HUM mice in early-stage drug efficacy studies of schistosomiasis. This strategy enables the prediction of pharmacokinetic exposures of medicines and late lead compounds that more closely reflect human metabolism, thereby allowing earlier and better-informed go/no-go decisions and minimising unnecessary medicinal chemistry optimisation driven by mouse-specific metabolic products.

The third phase of our investigation aimed to validate the 8HUM model as an improvement over WT mice in assessing the efficacy of PZQ against *S. mansoni*. In rac-PZQ dosed WT mice, *R*-PZQ and 4OH-PZQ exposures were higher in infected animals (compare Fig. 6D vs Fig. 5D); this difference is correlated with decreased hepatic expression of *cytochrome p450*s (Fig. 3A) and has been observed before (*36*). While we have not performed similar comparisons in 8HUM animals (uninfected vs infected, treated with the same dose of rac-PZQ), it is possible that the decreased expression of the ‘humanised’ *cytochrome p450*s in response to infection (Fig. 3A and Data file S3) may be responsible for the levelling out of *R*-PZQ and 4OH-PZQ exposures observed in infected 8HUM animals (Fig. 6C). Nevertheless, rac-PZQ is metabolised extremely rapidly in WT mice (Fig. 5 and (*37*)), necessitating a single high oral dose of 400 mg/kg body weight to sustain systemic - and likely mesenteric - exposures of *R*-PZQ sufficient to achieve substantial worm burden reductions (>90%) (*30*). Optimal PZQ-mediated efficacy in WT mice also depends on an intact immune system (*38, 39*). Although 8HUM mice exhibit immune responses to *S. mansoni* infection comparable to those of WT mice, their humanised cytochrome P450 profile resulted in markedly enhanced rac-PZQ efficacy, achieving >90% reductions in worm burdens (>70% reductions in egg burdens) at single oral doses of 100–200 mg/kg bodyweight; these parasitological reductions exceeded (tissue eggs) or were comparable (worms) to the efficacy observed in WT mice dosed with 400 mg/kg bodyweight (Fig. 6). Notably, while a single 100 mg/kg bodyweight oral dose of rac-PZQ produces (at most) a 53% reduction in worm burden in WT mice (*7*), the approximately two-fold increase in efficacy observed in 8HUM mice at the same dose underscores the translational advantage conferred by humanised drug metabolism in this model.

The 8HUM model incorporates the major human cytochrome P450 enzymes responsible for most CYP-mediated drug metabolism. However, it notably lacks CYP2C19 and CYP2J2. These isoforms, together with CYP1A2, CYP2C9, CYP2D6 and CYP3A4 (all of which are present in 8HUM mice), contribute substantially to rac-PZQ metabolism in humans (*40–42*). Consequently, direct extrapolation of findings related to Phase I metabolism from the 8HUM model to humans should be made with caution especially when also considering pharmacogenetic variations responsible for PZQ metabolism in endemic populations (*43*). Moreover, 8HUM mice retain several murine CYP subfamilies (e.g., Cyp2a, Cyp2b, and Cyp2e), whose residual activity - alongside that of Phase II conjugating enzymes and Phase III transporters - may influence the metabolic fate of novel compounds. Further humanisation of these CYP isoforms and Phase II/III proteins could enhance the translational fidelity of the 8HUM model for schistosomiasis drug discovery.

In summary, this study demonstrates that replacing WT mice with 8HUM animals in schistosome drug discovery pipelines effectively eliminates species-specific differences in drug metabolism, thereby aligning compound optimisation more closely with human pharmacokinetics. By enhancing translational relevance and improving the efficiency of development through reduced time and cost, 8HUM mice represent a powerful and practical refinement to WT mice, with the potential to transform preclinical evaluation and accelerate the discovery of human therapeutics for schistosomiasis.

## Materials and Methods

### Experimental Design

This study was designed to establish the 8HUM mouse model as a translational tool for assessing the *in vivo* efficacy of anti-schistosomal compounds during preclinical stages of drug discovery. The project had three phases. The first phase evaluated the 8HUM model as a suitable replacement for WT mice in supporting the sexual maturation of *S. mansoni* parasites and in permitting the development of infection-related characteristics. The second phase quantified the *in vitro* (intrinsic microsomal clearance, CL_int_) and *in vivo* (pharmacokinetics, PK) metabolism of racemic-praziquantel (rac-PZQ) in the 8HUM model. Using information derived from the PK studies, the third phase evaluated the efficacy of rac-PZQ in *S. mansoni* infected 8HUM mice at exposure levels far lower than that required to induce high levels (> 90%) of efficacy in WT mice. Experimental design was guided by power calculations (G*Power; https://www.gpower.hhu.de), previous experience and pilot studies. All animal work was undertaken in line with the 3R principles of replacement, reduction, and refinement (https://www.nc3rs.org.uk).

### Ethics

All regulated procedures described in this study were presented to and approved by the Animal Welfare and Ethical Review Bodies (AWERB) at Aberystwyth University (AU) and University of Dundee (UoD). They adhered to the United Kingdom Home Office Animals (Scientific Procedures) Act of 1986 and were performed according to project licenses PP2955700 (AU) and PP5016780 (UoD). Animals were inspected regularly by staff trained and experienced in small animal husbandry, with 24 hr access to veterinary advice. Mice were maintained with *ad libitum* access to food and water, and a twelve-hour light/dark period. Temperature and relative humidity were maintained between 20 °C and 24 °C, and 45% and 65%, respectively.

### Chemicals and reagents

Racemic-PZQ (rac-PZQ) for administration in PK studies was purchased from Sigma Aldrich/Merck (Burlington, MA, USA). (*R*)-PZQ, (*S*)-PZQ and 4OH-PZQ for bioanalysis were purchased from Stratech (Ely, UK), CliniSciences (Nanterre, France), and Accela ChemBio (San Diego, CA, USA), respectively. All LC–MS/MS mobile phase reagents were purchased from Fisher Scientific (Thermo Fisher Scientific, Waltham, MA, USA).

### *Schistosoma mansoni* lifecycle maintenance

For *S. mansoni* (Naval Medical Research Institute; NMRI strain) lifecycle maintenance, six- to twenty-six-week-old female TO (HsdOLa:TO - Tuck Ordinary; Envigo, UK) mice were percutaneously exposed by immersion of their tails in water containing 180 cercariae for 45 minutes. At 47 days post-infection, animals were administered an intra-peritoneal injection of a lethal dose of sodium pentobarbital containing 100 U/mL heparin and parasites collected by reverse perfusion of the hepatic-portal venous system. Following perfusion, eggs were recovered from infected livers, miracidia hatched and used to infect susceptible *Biomphalaria glabrata* snails (NMRI albino and pigmented outbred strains (*44*)).

### Phase 1 experimental infections

All mice included on study were housed in AU’s biological research facility for three weeks before investigations began. For experimental infection of transgenic 8HUM (*18*) (C57BL6/NTac background; PhaSER Biomedical, UK) and wild-type (WT) C57BL6/NCrl (Charles River Laboratories, UK) mice, six- to eight-week-old females (5-7 animals per group) were percutaneously exposed as above to 80 cercariae/mouse. Groups of uninfected animals (3 animals per group; 8HUM and WT) were housed alongside their infected counterparts. All mice on study were weighed individually, prior to, and weekly after experimental infection. At 46 days post-infection, blood samples (∼100 µL) were taken from the lateral tail veins of individual mice (infected and uninfected controls) for serum preparation. At 47 days post-infection, animals were sacrificed (as above) and murine-(spleens, livers, intestines, intestinal faecal pellets) and (where present; i.e. the infected animals) parasite-material (worms, liver eggs and intestinal eggs) obtained. These materials were processed for downstream parasitological, immunological, histological, pathological, transcriptional and microbiome analyses.

### Phase 2 Pharmacokinetic (PK) studies

Five uninfected female mice (2 x 8HUM and 3 x WT) were administered a single oral dose of rac-PZQ (25 mg/kg for 8HUM and 400 mg/kg for WT) as a fine suspension in 0.5% (w/v) hydroxypropyl methylcellulose (HPMC) at a dose volume of 5 mL/kg. Dose formulations were prepared on the day of dosing. Serial blood samples (10 μL per sample) were collected from the lateral tail vein prior to dosing, then at 5, 15, 30, 60, 120, 240, 360 and 480 min after administration. Each blood sample was diluted into 90 μL of Milli-Q ultrapure water, and stored at -20 °C prior to bioanalysis by U(H)PLC-MS/MS.

### Microsomal incubations

Microsome preparations and microsomal incubation procedures were described previously (*19*). Briefly, rac-PZQ was added at a final concentration of 0.5 µM to microsomes in buffer (0.5 mg/mL protein in 50 mM potassium phosphate, pH 7.4), and reactions initiated with addition of excess NADPH (final concentration 0.8 mg/mL). Aliquots of 50 µL were taken at 0, 3, 6, 9, 15 and 30 min, immediately mixed with two volumes of acetonitrile containing internal standard (IS, sulfadimethoxine, 50 ng/mL) and kept on ice. When all samples were collected, 250 µL of 20% acetonitrile was added to each and these were centrifuged for 10 min at 3,000 × *g* at ambient temperature. Supernatants were analysed by LC-MS/MS immediately.

### LC-MS/MS analysis

Samples from PK studies were analysed on an Acquity UPLC system coupled to a Xevo TQ-XS, operated using MassLynx software version 4.2 (Waters, Wilmslow, UK). Chromatographic separation was achieved using a Lux Cellulose-2 column, 150 × 4.6 mm, particle size 3 µm (part number 00F-4456-E0, Phenomenex, Macclesfield, UK). The column was held at 40 °C, mobile phase A was 20 mM ammonium formate in LC-MS grade water, mobile phase B was acetonitrile, flow rate was 1.2 mL/min, and the gradient programme was as follows: 0.0 – 1.0 min: 70% B, 1.0 – 2.0 min: 70 to 90% B, 2.0 to 9.5 min: 90% B, 9.5 – 10.0 min: 95 – 0% B, 10.0 – 10.5 min: 0 – 70% B and 10.5 – 11.5 min: 70% B. The mass spectrometer was operated with electrospray ionisation in positive mode, with capillary voltage of 0.8 kV, desolvation temperature of 600 °C, desolvation gas flow of 1000 L/h, cone gas flow of 150 L/h and source temperature of 150 °C. Multiple reaction monitoring (MRM) transitions were 313.2 > 203.1 for PZQ and 329.1 > 203.1 for 4OH-PZQ. Where samples containing rac-PZQ were analysed, only the *R* enantiomer (first of two peaks) was quantified. For 4OH-PZQ, two peaks were also observed due to the chiral chromatographic method used. Although it was considered likely that the first peak conformed to the *R* enantiomer and the second peak conformed to the *S* enantiomer (similar to what was observed for rac-PZQ), this could not be confirmed in the absence of authentic standards. Therefore, integrated areas of both peaks were summarised for reporting the total 4OH-PZQ levels.

Microsomal incubation samples were analysed on an Acquity UPLC system coupled to a Xevo TQS-Micro. Chromatographic separation was achieved using an Acquity UPLC BEH C18 column, 50 × 2.1 mm, particle size 1.7 µm (Waters, part number 186002350) held at 45 °C, with mobile phase A of 0.01% formic acid in Milli-Q water, and mobile phase B of 0.01% formic acid in LC-MS grade methanol. The gradient programme was as follows: 0.0 – 0.3 min: 5% B, 0.3 – 1.3 min: 5 to 95% B, 1.3 – 1.8 min: 95% B and 1.8 – 1.81 min: 95 – 5% B. Mass spectrometer parameters were as for PK sample analysis, with the exception of desolvation temperature which was set to 500 °C and cone gas flow which was 15 L/h. *R*- and *S*-enantiomers were not separated using this methodology and results are, therefore, reported as intrinsic clearance of racemic praziquantel.

### Intrinsic clearance determination

Raw LC-MS/MS data were exported to XLfit (IDBS, Woking, UK) for calculation of exponential decay rate constant (*k*) from the ratio of peak area of test compound to internal standard at each timepoint. Intrinsic clearance (CL_int_) was calculated by multiplying *k* by the incubation volume and the microsomal protein concentration. Verapamil was used as a routine positive control to confirm acceptable assay performance, in comparison to previous analytical runs. Lower and uppers limits of quantitation were 0.011 and 0.1 mL/min/mg, respectively. Data were scaled to mL/min/g (liver) using the scaling factors of 48 and 39.7 mg microsomal protein per g of liver for mouse and human, respectively.

### Quantitative bioanalysis of PK samples

On the day of sample analysis, calibration standards (CS) and quality controls (QC) were prepared from separate compound aliquots. Compounds were dissolved in DMSO and spiked into diluted blank blood as control matrix, prior to extraction with three volumes of acetonitrile containing internal standard (IS, sulfadimethoxine at 100 ng/mL), in parallel with PK samples. Concentration of drug in samples was determined by interpolation onto CS values using TargetLynx (Waters). Acceptance criteria of the bioanalytical method included CS accuracy of ± 20% of nominal (theoretical) concentration across the range of interest. Lower limit of quantification (LLoQ) was determined as the analyte response with ≥ three times the analyte response of single blanks (extracted blood containing IS but no test compound). QCs at low, medium, and high concentration levels were injected throughout the analytical run, with an acceptance criterion of 66% of injected samples being within ± 20% of nominal concentrations. All single and double blanks (extracted blood with no IS or test compound) were verified as free of interference at the retention times of the IS and test compound. Sample carryover was determined in a blank injection immediately following the injection of the top CS. Carryover was deemed accepted when the test compound response in the blank was < 1% of that in the top calibration sample. PK parameters were calculated by Non-Compartmental Analysis using Phoenix WinNonlin (Certara, Sheffield, UK).

### Phase 3 combined PK/Efficacy studies

All mice included on study were housed in AU’s biological research facility for four weeks before investigations began. For combined PK/efficacy studies, 8HUM and WT mice (six- to eight-week-old females) were percutaneously exposed to 120 cercariae/mouse. At 41 days post infection, groups of animals (9-10 animals per group) were administered, via oral gavage, a single dose (100 mg/kg and 200 mg/kg for 8HUM, 400 mg/kg for WT) of rac-PZQ as a fine suspension in 0.5% (w/v) HPMC at a dose volume of 5 mL/kg. Three animals from each dosing group were selected at random for PK evaluation and serial blood sampling of these animals was performed as outlined above. Infected animals (9-10 animals per group; both 8HUM and WT) treated with HPMC (5 mL/kg) only were included as negative controls for the efficacy experiment. At 47 days post-infection, animals were sacrificed (as above) and parasite material obtained and processed as before.

### Parasitology

Schistosomes were manually counted using a camera mounted to a Nexius Zoom dissection microscope (Euromex). Fresh livers and intestines (after removal of faecal pellets from the colon and flushing residual material with 10 mL PBS), were weighed and frozen at -80 °C until downstream processing. Faecal pellets were also stored at -80 °C until use. Livers and intestines were thawed and digested in 10 mL/g tissue 4% w/v potassium hydroxide (KOH) in ddH_2_O for 16 h at 37 °C with agitation as previously described (*45*). Eggs liberated from tissues were washed five times with 50 mL 0.01 M phosphate buffered saline (PBS), passed through a 100 µm cell strainer to remove residual debris and resuspended in 5 mL 4% (v/v) formaldehyde in ddH_2_O. Three x 10 µL aliquots per sample were diluted in water to a final volume of 200 µL, placed in a 96-well dark walled plate and imaged at 2.5x magnification using a Cytation 7 multi-modal imager (Agilent) with the GFP filter. Eggs were manually counted and eggs per gram of tissue (EPG) was calculated by extrapolating the mean number of eggs across three aliquots to the total volume of original sample and dividing by the tissue weight.

### Splenocyte culture and cytokine assays

Spleens were removed from each mouse, prior to reverse perfusion of the hepatic-portal venous system, and individual single cell suspensions prepared. Splenocytes were plated in 24-well tissue culture plates at a final concentration of 4 x 10^6^ cells/mL in complete media and cultured for 72 h in a humidified atmosphere containing 5% CO_2_ as previously described (*46*). Cells were stimulated with ConA (5 µg/mL), SEA (20 µg/mL), SWAP (50 µg/mL) or medium alone. SEA and SWAP were generated as described previously (*47, 48*) using mixed-sex adult worms as starting material (for SWAP). Supernatants were harvested at 72 h and assayed for IL-4, IL-5, IL-10, IFN-gamma, IL-13 and IL-17 by sandwich ELISA as previously described (*49*).

### Measurement of SWAP- and SEA-specific Ab responses

For assessment of anti-SEA and anti-SWAP Ig levels, serum samples prepared from whole blood collected at 46 days post-infection were assayed from individual animals as previously described with slight modifications (*46*). Briefly, individual mouse serum was serially diluted 1/100 - 1/102, 500 in 1% w/v BSA in PBS/0.05% Tween-20 and 50 µL was added to appropriate wells. Fifty µL of isotype specific horse-radish peroxidase (HRP) conjugated rabbit anti-mouse Abs (IgG_1_ = PA186329, Invitrogen; IgG_2b_ = 43R-IR032hrp_1MG, Fitzgerald Industries; total IgG = 616520, Invitrogen) in 1% w/v BSA in PBS/0.05% Tween-20 diluted at 1/1000 (measurement of IgG_1_ and IgG_2b_) or 1/2000 (measurement of total IgG) were added to the wells and incubated at 37 °C for 2 h. Reactions were developed at room temperature (RT) until the desired signal was reached (IgG_1_ = 14 – 21.5 min, IgG_2b_ = 22 – 36 min and Total IgG = 7.5 – 10.5 min), terminated with 100 µL of 1% w/v SDS and the OD (absorbance) at 405 nm was determined using a PolarStar Omega microtiter plate reader (BMG Labtech). Specific SWAP- and SEA-isotype titres were represented by the product of absorbance of a single point on the linear portion of the dilution curve.

### Liver histology

At day 47 post-infection, after reverse perfusion of the hepatic-portal venous system, approximately half of each liver was removed from individual mice, fixed in 10% buffered formalin for 24 h and then transferred to 70 % ethanol. Fixed liver samples were embedded in paraffin and 5 µm sections prepared for staining with haematoxylin and eosin (H&E) as well as picrosirius red (PSR) (*50*). To estimate percentage collagen within granuloma boundaries, PSR stained granulomas (counter stained with fast green for contrast) from 10x magnification micrographs captured using a Pannoramic 250 slide scanner were exported using 3D HISTECH Pannoramic Viewer (version 1.15.4). Granuloma boundaries were masked manually using the CVAT annotation service (www.cvat.ai). Annotations were exported in COCO 1.0 format (*51*), from which total areas (μm^2^), extrapolated spherical volumes (µm^3^) and percentage PSR stain (as a proxy for collagen) were calculated within object masks using a custom Python script (v. 3.10.12).

### Microbiome analyses of faecal samples

Genomic DNA was extracted from colonic content samples (day 47 post-infected- and uninfected mice) as well as no-template negative controls, using the PowerSoil DNA Isolation Kit (QIAGEN) following the manufacturer’s protocol. DNA library preparation and sequencing were performed using the Illumina MiSeq platform on the V3-V4 region with paired-end 250 bp reads. Raw 16S rRNA amplicon sequencing data are available from the European Nucleotide Archive (ENA) database under BioProject PRJEB94950.

All 16S rRNA sequencing datasets were initially processed using Cutadapt (version 4.1). Amplicon sequence analysis was conducted using the DADA2 pipeline (version 1.16). Quality filtering was performed with the filterAndTrim function, applying the following parameters: maxN = 0, maxEE = c(2, 2), truncQ = 2, rm.phix = TRUE, and truncLen = c(220, 200). Chimeric amplicon sequence variants (ASVs) were removed using the removeBimeraDenovo function with the ‘consensus’ method. Taxonomic assignment was performed by aligning ASVs to the SILVA database (version 138) curated for DADA2.

The resulting ASV and taxonomy tables were further analysed using MicrobiomeAnalyst (*52*). Data were normalised by cumulative sum scaling (CSS) and rarefied to a uniform sequencing depth of default reads per sample. Alpha diversity metrics, including species richness and Shannon diversity index, were compared between groups using unpaired and pairwise Kruskal-Wallis tests. Beta diversity was evaluated using nonmetric multidimensional scaling (NMDS) based on Bray–Curtis dissimilarity metrics. Statistical differences between groups were assessed by Permutational Multivariate Analysis of Variance (PERMANOVA).

### RNA-sequencing (RNA-Seq)

At 47 days post-infection, 250 mg of liver tissue/mouse was removed from the left lobe prior to reverse perfusion of the hepatic-portal venous system and total RNA isolated as previously described (*53*). Total RNA was also isolated from adult male and female worms at 47 days post-infection. Here, schistosomes from three individual mice (WT, *n* = 3 and 8HUM, *n* = 3) were pooled by sex and RNA isolated using the MasterPure Complete DNA and RNA Purification Kit (Cambio) following the manufacturer’s instructions but with a 60-minute DNase treatment. The quality and quantity of murine and schistosome total RNA samples were assessed via analysis on a ThermoFisher Nanodrop 2000 and an Agilent Bioanalyzer 2100. Total RNA samples were submitted to BGI Tech Solutions Co., Limited (Poland) for library construction and DNBSEQ eukaryotic strand-specific transcriptome resequencing at paired-end 100 bp read length with ≥22 million clean reads per sample. All RNA-Seq samples used in this study are available to download from ENA BioProject PRJEB94950.

### Transcriptomic assembly and analysis

All mouse samples were assembled against the *Mus musculus* reference genome (GRCm39, NCBI RefSeq accession GCF_000001635.27) by BGI Tech Solutions Co., Limited (Poland). Briefly, reads were mapped using HISAT (v.2.04) (*54*) followed by quantification of transcript expression using Bowtie2 (v.2.25) (*55*) and RSEM (v.1.2.8). To delineate human sequence reads from mouse reads for the replaced cytochromes and transcription factors, a chimeric HISAT2 reference was built to include both human and mouse cytochrome sequences in the GRCm39 core transcriptome. All samples (WT and 8HUM) were remapped against this reference for the purposes of cytochrome characterisation only (i.e., not for downstream analyses subsequently described). Differential expression was quantified using DESeq2 (*56*) with a minimal *q*-value cutoff of ≤ 0.05. Functional annotation of genes was performed for GO terms by mapping genes to the Gene Ontology database (http://www.geneontology.org/) followed by Gene Set Enrichment Analysis (GSEA) in the phyper R package.

Given that *S. mansoni* is a non-reference species, assembly and analysis was performed in-house at AU. Using version 10 (v.10) of the *S. mansoni* genome assembly on WormBase ParaSite (PRJE36577), a genome index was constructed using soft-masked sequences and mRNA transcript decoys in Salmon (k-mer length = 27) with collapsing of identical transcripts permitted (indexing transcripts for 9,980 protein coding genes). Transcript quantification was also performed using Salmon, followed by differential expression analysis by edgeR (*57*) in Trinity (v.2.15.2) with cut-offs set at *q* ≤ 0.001 and a minimum absolute log_2_FC of 2 (4-fold change).

Whole transcriptome heatmaps were generated using the heatmap.2 function of Bioconductor’s gplots with the RcolorBrewer palette library in R (v. 4.5.1) using Rstudio (v. 2024.12.0). Default parameters were used according to documentation, with row-means dendrogram clustering applied using Rowv = TRUE. Subset heatmaps and bubbleplots for GSEA analysis were generated in Python (v.3.10.12) by leveraging matplotlib in a custom script.

### Statistical analyses

Distributional assessments were performed prior to all statistical testing using GraphPad Prism (v.8.4.3). Parametric datasets were compared by either a one-way or two-way ANOVA with Tukey’s Multiple Comparisons Test. For non-parametric data, differences were assessed using a Mann-Whitney U test for two groups. All significance values given in this manuscript are adjusted as required and significant *p* or *q* values are abbreviated as *, **, *** and **** for < 0.05, < 0.01, < 0.001 and < 0.0001, respectively. All statistical comparisons and corrections for transcriptomic data were performed using default options within the respective differential expression software (edgeR for worm data and DESeq2 for mouse data).

## List of Supplementary Materials

**Fig. S1.**
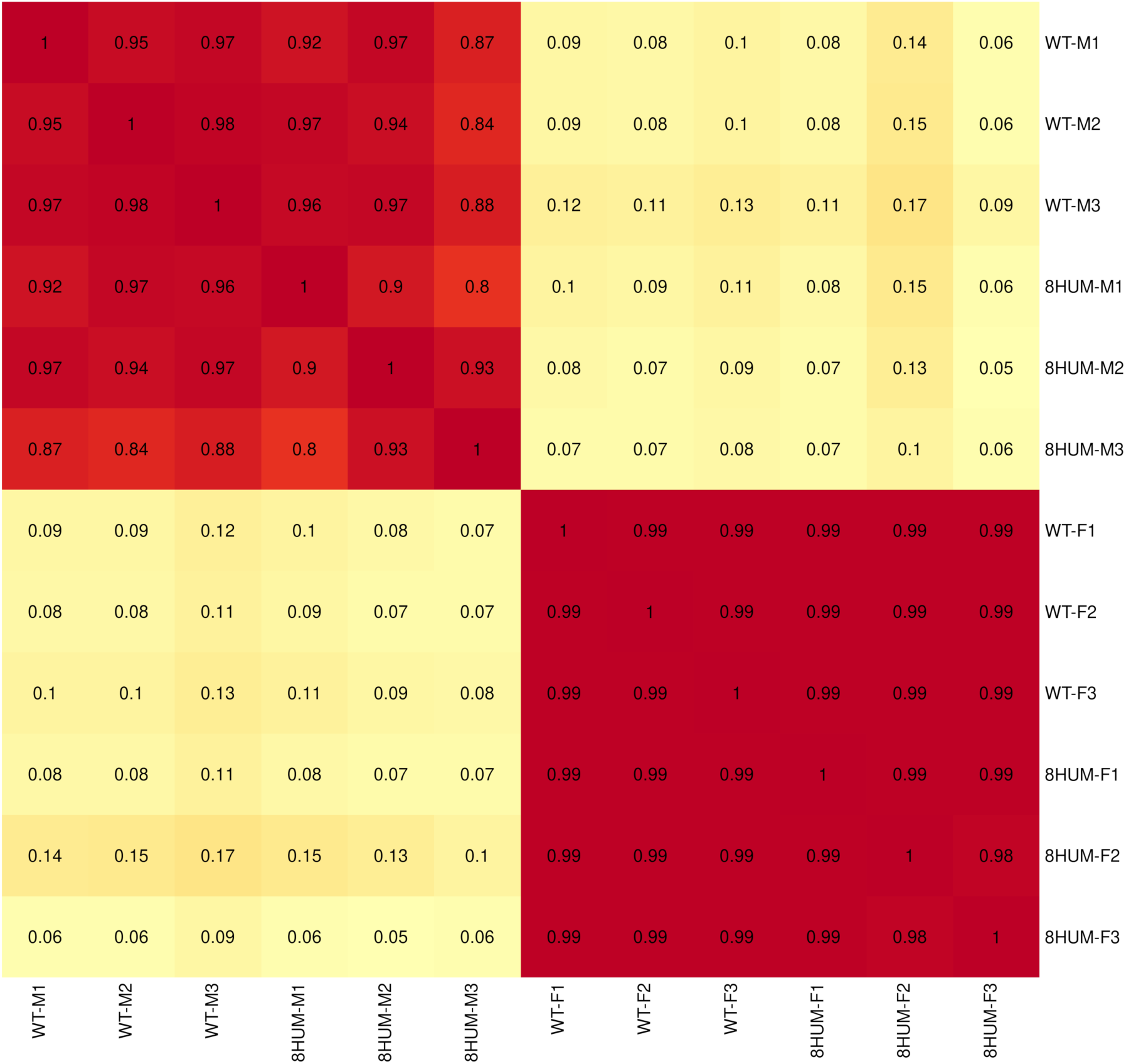
Pearson correlation matrix of transcript expression between samples for male (*n* = 3 per strain) and female (*n* = 3 per strain) worms derived from WT and 8HUM mice.

**Fig. S2.**
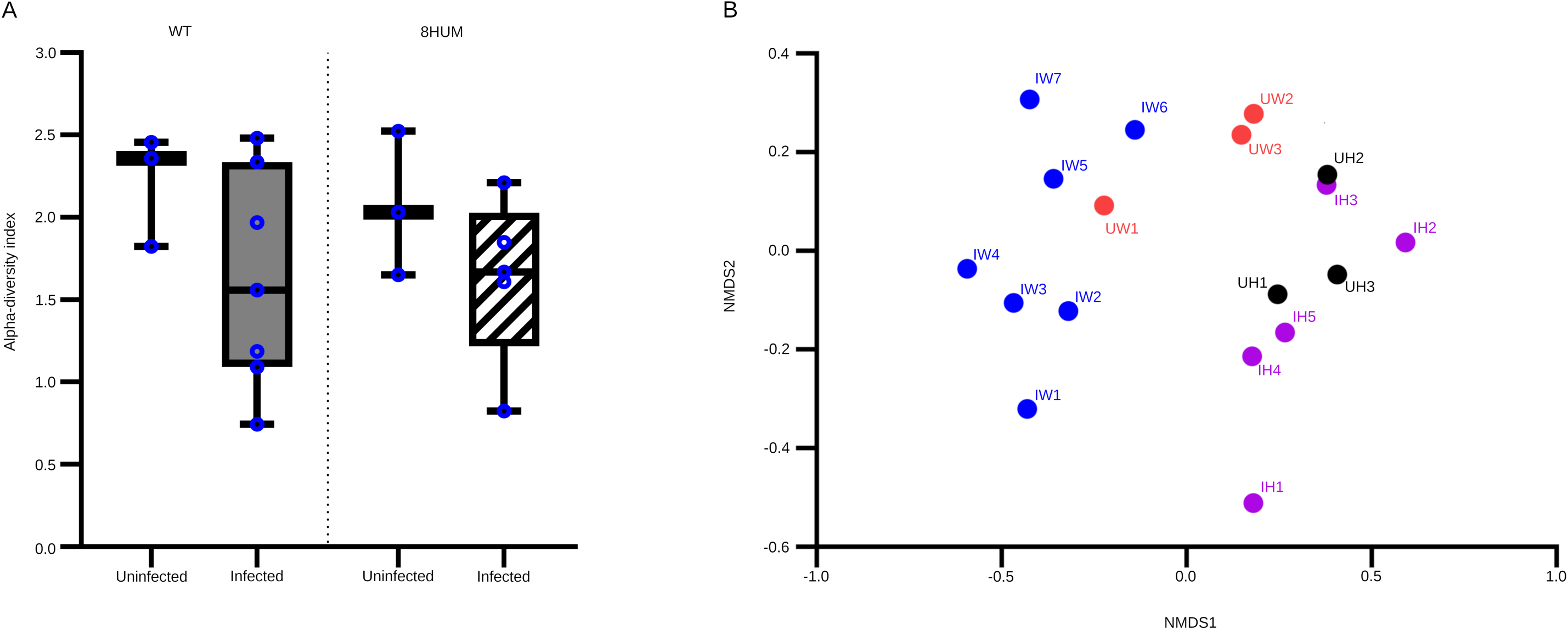
Compositional profiles of the faecal microbiomes for each mouse. **(A)** Alpha diversity using species richness and Shannon index (*p* value: 0.31284; Kruskal–Wallis statistic: 3.5617). **(B)** Beta diversity using non-metric multidimensional scaling (NMDS) (PERMANOVA: F-value: 1.9469; R²: 0.29438; *p* value: 0.112; NMDS Stress = 0.12384). WT uninfected: UW1-3 (red), WT infected: IW1-7 (blue), 8HUM uninfected: UH1-3 (black), and 8HUM infected: IH1-5 (purple).

**Fig. S3.**
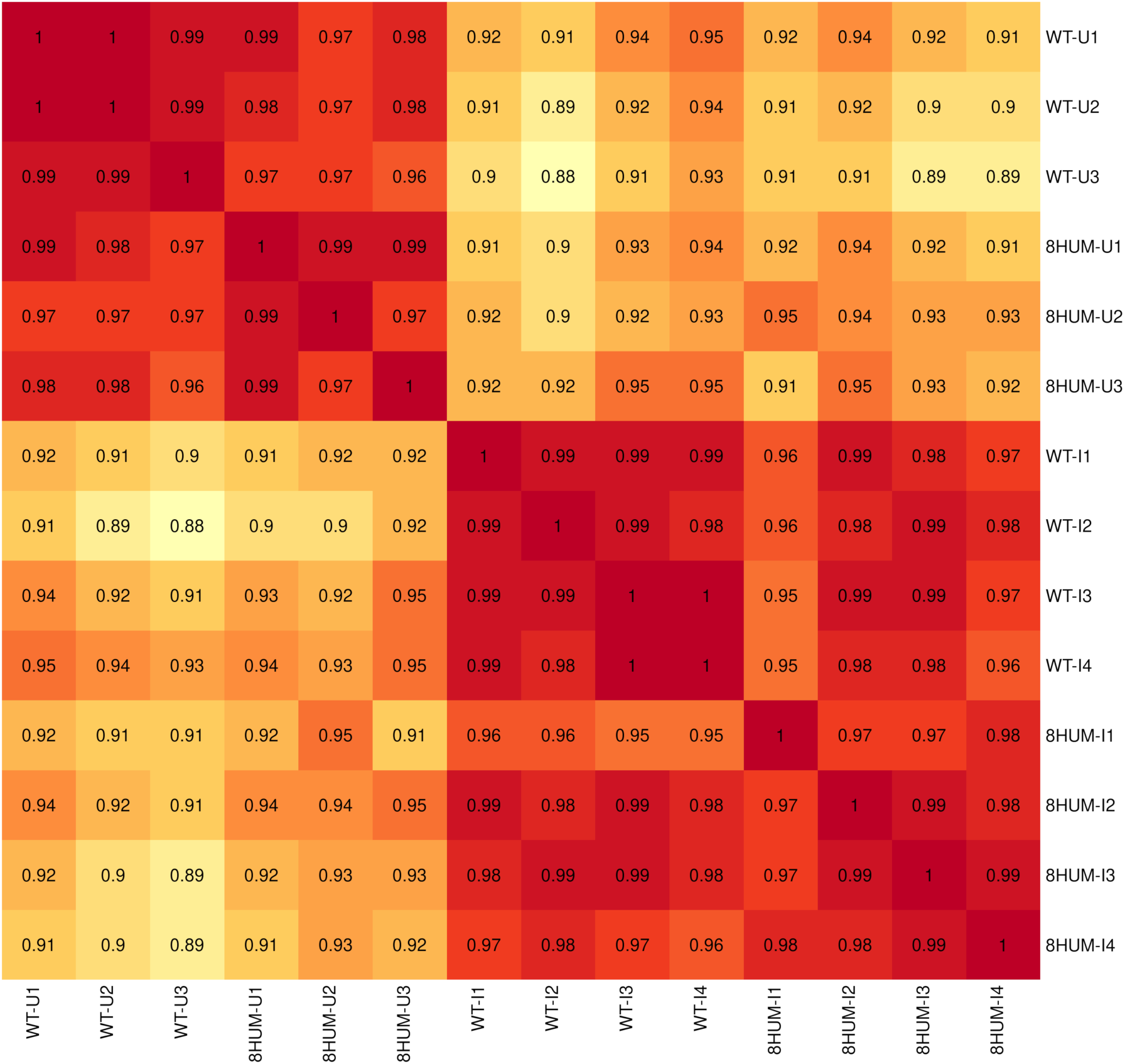
Pearson correlation matrix of gene expression between liver samples for uninfected (*n* = 3 per strain) and *S. mansoni* infected (*n* = 4 per strain) wild-type (WT) and 8HUM mice.

**Fig. S4.**
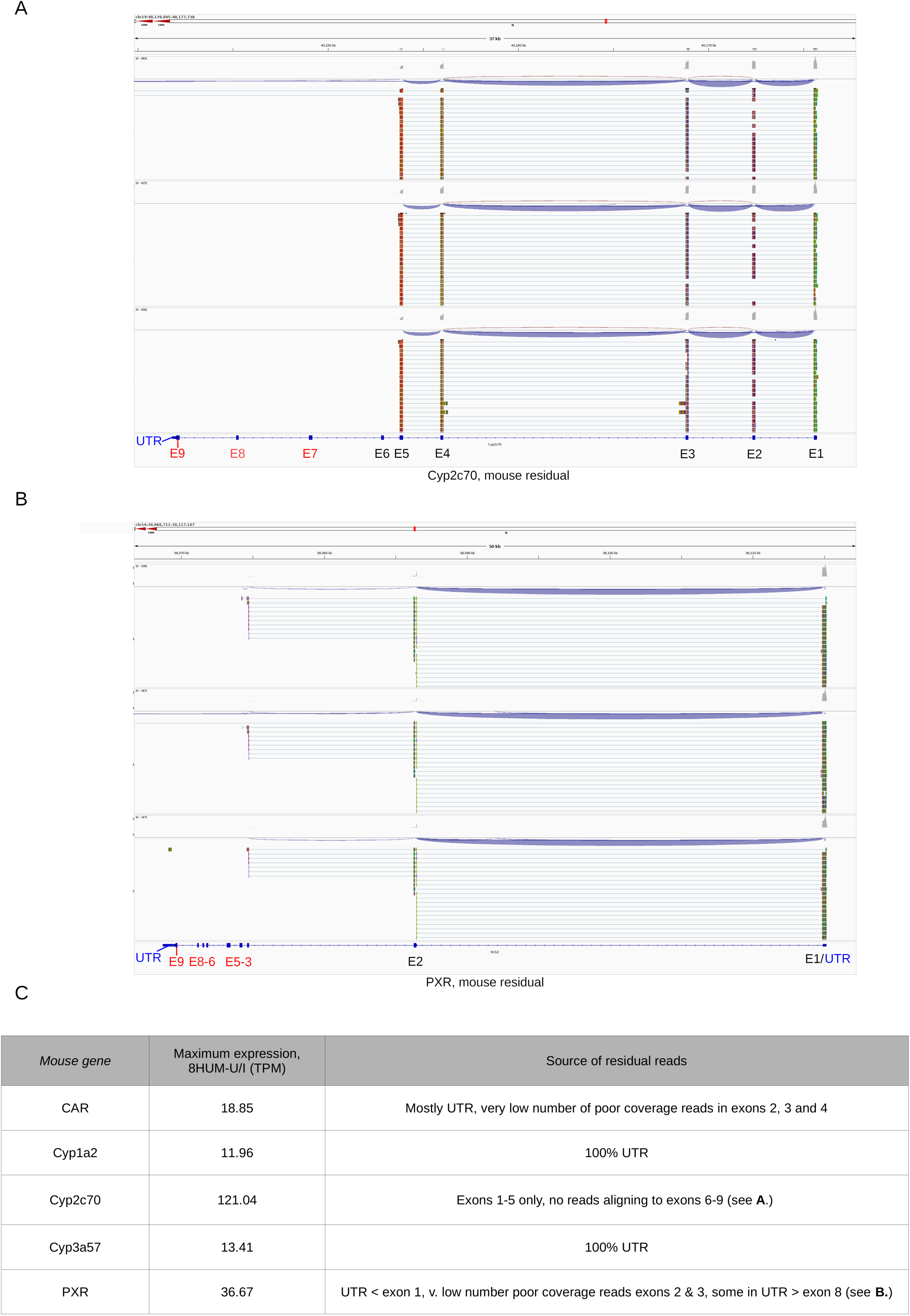
IGV interrogation of residual mouse reads mapping to 8HUM samples following removal of reads mapping to humanised genes (panels A and B). Where mouse genes were detectable in 8HUM samples at >10 TPM, explanations for residual reads are provided in Table C.

Data file S1 log_2_ transformed transcript per million (TPM) counts for 9,890 *S. mansoni* coding genes in male (-M) and female (-F) worms derived from wild-type (WT) and 8HUM mice.

Data file S2 Differentially expressed transcripts for male and female worms between WT and 8HUM mice at *q* < 0.001, 0.01 and 0.05 and log_2_FC > 2 (FC > 4).

Data file S3 log2 transformed transcript per million (TPM) expression of the 35 mouse cytochromes and 2 transcription factors substituted for human orthologues in 8HUM mice as well as their humanised replacements. Gene expression is given for both uninfected (*n* = 3 per strain) and *S. mansoni* infected (*n* = 4 per strain) individuals. Mapped using chimeric HISAT2 reference.

Data file S4 log_2_ transformed transcript per million (TPM) counts for 17,572 mouse protein coding genes in uninfected and *S. mansoni* infected wild-type (WT) and 8HUM mice hepatic transcriptomes.

Data file S5 Differentially expressed genes in wild-type (WT) and 8HUM mice hepatic transcriptomes in response to *S. mansoni* infection.

Data file S6 Top twenty enriched GO term gene sets for Biological Processes (BP) in wild-type (WT) and 8HUM mice hepatic transcriptomes in response to *S. mansoni* infection.

Data file S7 Top twenty enriched GO term gene sets for and Molecular Function (MF) in wild-type (WT) and 8HUM mice hepatic transcriptomes in response to *S. mansoni* infection.

Data file S8 log_2_ transformed transcript per million (TPM) counts and enrichment data for genes categorised under granuloma-associated GO term collagen fibril organisation (GO:0030199) in wild-type (WT) and 8HUM mice hepatic transcriptomes.

Data file S9 log2 transformed transcript per million (TPM) counts and enrichment data for genes categorised under granuloma-associated GO term extracellular matrix organisation (GO:0030198) in wild-type (WT) and 8HUM mice hepatic transcriptomes.

Reference (*58*) cited in Data file S2 for spatial mapping data.

## Acknowledgements

We acknowledge Ms Julie Hirst, Mr Rory Geoghegan and Dr Graham Brand for assisting in the maintenance of the schistosome lifecycle. We thank Ms Charly Morgan for assisting with the 16S rRNA work (library preparation and DNA sequencing).

## Author contributions

Conceptualisation: KFH, KDR

Methodology: BJH, SDD, JEF-T, KL, AHC, ME, GR, AKM, LF, LS, FRCS, AT, OP, YHL, MTS, WA, MH

Investigation: KFH, BJH, SDD, JEF-T, KL, AHC, ME, GR, LS, FRCS, RP, IWC, BD, BKO, VHG

Visualisation: KFH, BJH, SDD, JEF-T, KL, AHC, AKM, LF, AT

Funding acquisition: KFH, IHG, BB, KDR, GR

Project administration: KFH, ASM, BB, NC, KDR, IHG Software: SDD

Data curation: SDD

Supervision: KFH, MTS, WA, BB, KDR, IHG

Writing – original draft: KFH, BJH, SDD, JEF-T, KL, AKM, YHL

Writing – review & editing: KFH, BJH, SDD, JEF-T, KL, AHC, ME, GR, AKM, RP, OP, YHL, IWC, BD, BKO, VHG, MH, MTS, WA, ASM, BB, NC, KDR, IHG

## Contributing interests

Authors declare that they have no competing interests.

## Data and materials availability

All data are available in the main text, accession numbers or the supplementary materials.

## References

1. D. P. McManus et al., Schistosomiasis. Nat Rev Dis Primers 4, 13 (2018).

2. S. K. Park et al., Mechanism of praziquantel action at a parasitic flatworm ion channel. Sci Transl Med 13, eabj5832 (2021).

3. W. Le Clec’h et al., Genetic analysis of praziquantel response in schistosome parasites implicates a transient receptor potential channel. Sci Transl Med 13, eabj9114 (2021).

4. J. D. Chan et al., The anthelmintic praziquantel is a human serotoninergic G-protein-coupled receptor ligand. Nat Commun 8, 1910 (2017).

5. N. Vale et al., Praziquantel for Schistosomiasis: Single-drug metabolism revisited, mode of action, and resistance. Antimicrob Agents Chemother 61, (2017).

6. P. Olliaro, P. Delgado-Romero, J. Keiser, The little we know about the pharmacokinetics and pharmacodynamics of praziquantel (racemate and R-enantiomer). J Antimicrob Chemother 69, 863–870 (2014).

7. R. Gonnert, P. Andrews, Praziquantel, a new board-spectrum antischistosomal agent. Z Parasitenkd 52, 129–150 (1977).

8. R. E. Sanya et al., The impact of intensive versus standard anthelminthic treatment on allergy-related outcomes, helminth infection intensity and helminth-related morbidity in Lake Victoria fishing communities, Uganda: results from the LaVIISWA cluster randomised trial. Clin Infect Dis, (2018).

9. T. Crellen et al., Reduced efficacy of praziquantel against *Schistosoma mansoni* is associated with multiple rounds of mass drug administration. Clin Infect Dis 63, 1151–1159 (2016).

10. D. J. Berger et al., Extensive transmission and variation in a functional receptor for praziquantel resistance in endemic *Schistosoma mansoni*. bioRxiv, (2024).

11. WHO, D. o. c. o. n. t. diseases, Ed. (WHO, http://www.who.int/neglected_diseases/resources/WHO_HTM_NTD_2012.1/en/, 2012).

12. N. Caldwell et al., Perspective on Schistosomiasis drug discovery: Highlights from a Schistosomiasis drug discovery workshop at Wellcome Collection, London, September 2022. ACS Infect Dis 9, 1046–1055 (2023).

13. W. E. Evans, M. V. Relling, Pharmacogenomics: translating functional genomics into rational therapeutics. Science 286, 487–491 (1999).

14. L. C. Wienkers, T. G. Heath, Predicting *in vivo* drug interactions from *in vitro* drug discovery data. Nat Rev Drug Discov 4, 825–833 (2005).

15. A. Saravanakumar, A. Sadighi, R. Ryu, F. Akhlaghi, Physicochemical properties, biotransformation, and transport pathways of established and newly approved medications: A systematic review of the top 200 most prescribed drugs vs. the FDA-approved drugs between 2005 and 2016. Clin Pharmacokinet 58, 1281–1294 (2019).

16. M. A. Cerny, Prevalence of non-cytochrome P450-mediated metabolism in food and drug administration-approved oral and intravenous drugs: 2006-2015. Drug Metab Dispos 44, 1246–1252 (2016).

17. D. R. Nelson et al., Comparison of cytochrome P450 (CYP) genes from the mouse and human genomes, including nomenclature recommendations for genes, pseudogenes and alternative-splice variants. Pharmacogenetics 14, 1–18 (2004).

18. C. J. Henderson et al., An extensively humanized mouse model to predict pathways of drug disposition and drug/drug interactions, and to facilitate design of clinical trials. Drug Metab Dispos 47, 601–615 (2019).

19. A. K. MacLeod et al., Acceleration of infectious disease drug discovery and development using a humanized model of drug metabolism. Proc Natl Acad Sci U S A 121, e2315069121 (2024).

20. J. D. Liberatos, *Schistosoma mansoni*: male-biased sex ratios in snails and mice. Exp Parasitol 64, 165–177 (1987).

21. A. Cortes et al., Baseline gut microbiota composition is associated with *Schistosoma mansoni* infection burden in rodent models. Front Immunol 11, 593838 (2020).

22. T. P. Jenkins et al., *Schistosoma mansoni* infection is associated with quantitative and qualitative modifications of the mammalian intestinal microbiota. Sci Rep 8, 12072 (2018).

23. K. F. Hoffmann et al., Disease fingerprinting with cDNA microarrays reveals distinct gene expression profiles in lethal type 1 and type 2 cytokine-mediated inflammatory reactions. Faseb J 15, 2545–2547. (2001).

24. A. H. Costain et al., Corrigendum: Dynamics of host immune response development during *Schistosoma mansoni* infection. Front Immunol 14, 1229665 (2023).

25. K. F. Hoffmann, T. A. Wynn, D. W. Dunne, Cytokine-mediated host responses during schistosome infections; walking the fine line between immunological control and immunopathology. Adv Parasitol 52, 265–307 (2002).

26. M. G. Chiaramonte, D. D. Donaldson, A. W. Cheever, T. A. Wynn, An IL-13 inhibitor blocks the development of hepatic fibrosis during a T-helper type 2-dominated inflammatory response. J Clin Invest 104, 777–785 (1999).

27. S. K. Madala et al., Matrix metalloproteinase 12-deficiency augments extracellular matrix degrading metalloproteinases and attenuates IL-13-dependent fibrosis. Journal of immunology 184, 3955–3963 (2010).

28. B. Vaillant, M. G. Chiaramonte, A. W. Cheever, P. D. Soloway, T. A. Wynn, Regulation of hepatic fibrosis and extracellular matrix genes by the th response: new insight into the role of tissue inhibitors of matrix metalloproteinases. Journal of immunology 167, 7017–7026 (2001).

29. C. R. Perry et al., Differential expression of chemokine and matrix re-modelling genes is associated with contrasting schistosome-induced hepatopathology in murine models. PLoS Negl Trop Dis 5, e1178 (2011).

30. I. Meister et al., Activity of praziquantel enantiomers and main metabolites against *Schistosoma mansoni*. Antimicrob Agents Chemother 58, 5466–5472 (2014).

31. G. Yamey et al., Vaccine value profile for schistosomiasis. Vaccine 64, 126020 (2025).

32. J. M. F. Gardner, N. R. Mansour, A. S. Bell, H. Helmby, Q. Bickle, The discovery of a novel series of compounds with single-dose efficacy against juvenile and adult *Schistosoma* species. PLoS Negl Trop Dis 15, e0009490 (2021).

33. S. Kaojarern, S. Nathakarnkikool, U. Suvanakoot, Comparative bioavailability of praziquantel tablets. DICP 23, 29–32 (1989).

34. A. Metwally, J. Bennett, S. Botros, F. Ebeid, D. el attar Gel, Impact of drug dosage and brand on bioavailability and efficacy of praziquantel. Pharmacol Res 31, 53–59 (1995).

35. G. Zdesenko, F. Mutapi, Drug metabolism and pharmacokinetics of praziquantel: A review of variable drug exposure during schistosomiasis treatment in human hosts and experimental models. PLoS Negl Trop Dis 14, e0008649 (2020).

36. M. A. Gotardo et al., Modulation of expression and activity of cytochrome P450s and alteration of praziquantel kinetics during murine schistosomiasis. Mem Inst Oswaldo Cruz 106, 212–219 (2011).

37. N. Abla et al., Evaluation of the pharmacokinetic-pharmacodynamic relationship of praziquantel in the *Schistosoma mansoni* mouse model. PLoS Negl Trop Dis 11, e0005942 (2017).

38. P. J. Brindley, A. Sher, The chemotherapeutic effect of praziquantel against *Schistosoma mansoni* is dependent on host antibody response. J Immunol 139, 215–220 (1987).

39. J. Modha, J. R. Lambertucci, M. J. Doenhoff, D. J. McLaren, Immune dependence of schistosomicidal chemotherapy: an ultrastructural study of *Schistosoma mansoni* adult worms exposed to praziquantel and immune serum in vivo. Parasite Immunol 12, 321–334 (1990).

40. X. Q. Li, A. Bjorkman, T. B. Andersson, L. L. Gustafsson, C. M. Masimirembwa, Identification of human cytochrome P(450)s that metabolise anti-parasitic drugs and predictions of *in vivo* drug hepatic clearance from in vitro data. Eur J Clin Pharmacol 59, 429–442 (2003).

41. H. Wang et al., Metabolic profiling of praziquantel enantiomers. Biochem Pharmacol 90, 166–178 (2014).

42. T. D. Gebreyesus et al., CYP2C19 and CYP2J2 genotypes predict praziquantel plasma exposure among Ethiopian school-aged children. Sci Rep 14, 11730 (2024).

43. R. H. Mnkugwe, O. Minzi, S. Kinung’hi, A. Kamuhabwa, E. Aklillu, Effect of pharmacogenetics variations on praziquantel plasma concentrations and Schistosomiasis treatment outcomes among infected school-aged children in Tanzania. Front Pharmacol 12, 712084 (2021).

44. K. K. Geyer et al., The *Biomphalaria glabrata* DNA methylation machinery displays spatial tissue expression, is differentially active in distinct snail populations and is modulated by interactions with *Schistosoma mansoni*. PLoS Negl Trop Dis 11, e0005246 (2017).

45. A. W. Cheever, Conditions affecting the accuracy of potassium hydroxide digestion techniques for counting *Schistosoma mansoni* eggs in tissues. Bull World Health Organ 39, 328–331 (1968).

46. K. F. Hoffmann, S. L. James, A. W. Cheever, T. A. Wynn, Studies with double cytokine-deficient mice reveal that highly polarized Th1- and Th2-type cytokine and antibody responses contribute equally to vaccine-induced immunity to *Schistosoma mansoni*. J Immunol 163, 927–938. (1999).

47. E. J. Pearce et al., *Schistosoma mansoni* in IL-4-deficient mice. Int Immunol 8, 435–444 (1996).

48. A. T. Vella, E. J. Pearce, CD4^+^ Th2 response induced by *Schistosoma mansoni* eggs develops rapidly, through an early, transient, Th0-like stage. J.Immunol. 148, 2283–2288 (1992).

49. A. H. Costain et al., Dynamics of Host Immune Response Development During *Schistosoma mansoni* Infection. Front Immunol 13, 906338 (2022).

50. K. Su et al., SOX9 plays an essential role in myofibroblast driven hepatic granuloma integrity and parenchymal repair during schistosomiasis-induced liver damage. PLoS Pathog 21, e1012928 (2025).

51. T.-Y. Lin et al., in *Computer Vision–ECCV 2014: 13th European Conference, Zurich, Switzerland, September 6-12*, 2014, Proceedings, Part V 13. (Springer, 2014), pp. 740–755.

52. A. Dhariwal et al., MicrobiomeAnalyst: a web-based tool for comprehensive statistical, visual and meta-analysis of microbiome data. Nucleic Acids Res 45, W180–W188 (2017).

53. K. F. Hoffmann, J. M. Fitzpatrick, Gene expression studies using self-fabricated parasite cDNA microarrays. Methods Mol Biol 270, 219–236 (2004).

54. D. Kim, B. Langmead, S. L. Salzberg, HISAT: a fast spliced aligner with low memory requirements. Nat Methods 12, 357–360 (2015).

55. B. Langmead, S. L. Salzberg, Fast gapped-read alignment with Bowtie 2. Nat Methods 9, 357–359 (2012).

56. M. I. Love, W. Huber, S. Anders, Moderated estimation of fold change and dispersion for RNA-seq data with DESeq2. Genome biology 15, 550 (2014).

57. M. D. Robinson, D. J. McCarthy, G. K. Smyth, edgeR: a Bioconductor package for differential expression analysis of digital gene expression data. Bioinformatics 26, 139–140 (2010).

58. G. Wendt et al., A single-cell RNA-seq atlas of *Schistosoma mansoni* identifies a key regulator of blood feeding. Science 369, 1644–1649 (2020).

